# MAFB and MAF cooperatively orchestrate human anti-inflammatory macrophage identity via autocrine Interleukin-10

**DOI:** 10.64898/2026.11.11.765660

**Authors:** Alejandro del Castillo-Buey, Israel Ríos, Miriam Simón-Fuentes, Alicia Capel-Jorquera, Cristina Herrero, Ángel Maldonado, Inés Zapico, Felipe Clemente, Alba Angelina, Oscar Palomares, Miguel A. Vega, Ángel L. Corbí

## Abstract

Macrophage lineage identity is governed by intricate transcriptional networks. Although the large MAF basic region-leucine zipper (bZIP) transcription factors MAFB and MAF are critical regulators of anti-inflammatory programs, their molecular coordination and non-redundant contributions within human macrophages remain poorly defined. Here, utilizing integrated proteomic, transcriptomic, and epigenomic profiling, we demonstrate that endogenous MAFB and MAF physically associate to form a stable heterodimeric complex on chromatin in human M-CSF-dependent monocyte-derived macrophages (M-MØ). Genome-wide CUT&Tag mapping reveals a hierarchical epigenomic architecture in which MAFB acts as the primary global anchor of M-MØ identity, whereas MAF provides localized transcriptional tuning. Mechanistically, both factors directly bind the *IL10* locus, but functional IL-10 expression is primarily driven by MAFB. Neutralizing antibody assays reveal that endogenously produced IL-10 establishes an obligate autocrine feedback loop mandatory for both the initial acquisition and the maintenance of the M-MØ anti-inflammatory profile. Functionally, MAFB and MAF exert non-redundant control over core macrophage effectors: MAFB dictates apoptotic cell efferocytosis amd T cell activation, whereas MAF modulates phagocytosis, and acts as a brake suppressing acute pro-inflammatory cytokine/chemokine hyper-induction. Together, these findings establish a comprehensive molecular blueprint of how MAFB and MAF cooperatively fine-tune human monocyte-derived macrophage identity and functional plasticity.

## INTRODUCTION

Tissue-resident macrophages (TRM) are pivotal custodians of tissue homeostasis that orchestrate organogenesis and fine-tune immune responses through efferocytosis, phagocytosis, pathogen recognition, and tissue repair ^1^. Besides these core functions, TRM exhibit remarkable functional plasticity as they adapt their transcriptomic identity to specific anatomical microenvironments ^2^. Functional specialization of TRMs is dictated not only by local cues but also by their developmental origin (embryonically-derived *vs.* monocyte-derived) ^1^. Macrophage Colony-Stimulating Factor (M-CSF) is essential for the generation, maintenance and self-renewal of TRM ^3^, with most M-CSF activities being executed primarily through induction of the basic region-leucine zipper (bZIP) transcription factors MAFB and MAF ^4,5^. Combined disruption of MAFB and MAF causes primary macrophages to proliferate exponentially in response to M-CSF stimulation ^6,7^ by releasing the constraints on Ets-1/2 and PU.1 and activating c-Myc and KLF4 expression ^4,8,9^, and induces proliferation and a proinflammatory/anti-tumoral reprogramming in Kupffer cells ^10^. Although MAFB and MAF exhibit distinct expression pattern in macrophages ^11^, they contribute to establishing macrophage identity and restricting self-renewal ^3,6^. Indeed, the M-CSF-dependent phenotypic and functional specialization of human monocyte-derived macrophages (M-MØ) heavily relies on MAFB and MAF ativity ^12−17^.

MAFB is a key driver of genes essential for TRM functions and lineage commitment ^18^, influences actin-dependent macrophage morphology ^19^, and profoundly shapes the anti-inflammatory (IL-10-producing), reparative and immunosuppressive properties of M-MØ ^11−1^^6,20^. MAFB promotes the expression of anti-inflammatory genes like *ADGRE1* ^21^ and *IL10* ^22,23^, upregulates ABCA1/ABCG1 cholesterol transporters ^23^, enhances the production of specialized pro-resolving mediators ^24^, and dampens type I interferon (IFN) production ^25^ by blocking coactivator recruitment to IRF3 ^26^. The coexpression of MAFB and its target genes in human CD163^+^ TRM and tumor-associated macrophages (TAM) strongly supports its role in maintaining homeostatic and immunosuppressive functions ^22^, with MAFB downregulation shifting the macrophage phenotype toward a pro-inflammatory state ^22^. Similarly, MAF regulates myeloid self-renewal ^6^, shapes macrophage identity through expression of markers including CD206, CD13 (*ANPEP*) ^27^, F4/80 (*ADGRE1*) ^21,28^ and CSF1R ^29^, and favors an anti-inflammatory profile by positively regulates IL-10 expression ^29−3^^3^.

MAF also shapes the immunosuppressive transcriptome and functions of TAM ^29,34,35^, and acts as a metabolic checkpoint regulating the TCA cycle and UDP-GlcNAc biosynthesis to favor M2-like macrophage activation ^29^. Moreover, MAF is transiently downregulated during acute inflammation, allowing macrophages to temporarily adopt a pro-inflammatory state before re-expressing MAF to enforce inflammation resolution ^36^.

Despite their overlaping roles roles, MAFB and MAF also exhibit distinct, context-specific functions. A high MAFB/MAF expression ratio in lung macrophages correlates with exacerbated pro-fibrotic responses in severe COVID-19 ^37,38^. In lung development, MAFB dictates monocyte differentiation into mouse interstitial macrophages (IM), whereas MAF selectively controls CD206+ IM subset identity ^18^. MAF also plays an essential role in perivascular macrophages regulating the metabolic syndrome ^39^ and even protection against Alzheimer’s disease ^40^. These MAFB-and MAF-specific effects resemble their distinct effects on the differentiation of auditory neurons ^41^ and during maturation of postnatal cortical interneuron ^42^. The differential effects of MAFB and MAF could be partly explained by their distinct dimerization and interacting patners. *In vitro*-synthesized MAFB heterodimerizes with MAF and FOS, but not with JUN ^17^, whereas *in vitro*-synthesized MAF binds FOS and JUN, and its DNA-binding ability is inhibited by USF2 ^27^. Over-expression and *in vitro* experiments indicate that MAFB interacts with IRF-7 ^25,26^, Spi-B ^25^, Ets1 ^43^ and MYB ^44,45^, whereas MAF binds MYB and Ets1 ^45^. Regarding endogenous proteins, MAF and MAFB physically interact in differentiated keratinocytes ^46^, MAFB associates with IRF3 in HeLa and HepG2 cells ^26^, and MAF interacts with AhR in T lymphocytes ^47^. However, the direct physical interaction of endogenous MAFB and MAF proteins and their precise division of labor in human monocyte-derived macrophages remain unresolved. Here, we integrate proteomic, transcriptomic, and genome-wide epigenomic profiling in M-MØ to define the co-regulated and factor-specific cistromic networks governed by MAFB and MAF. We reveal that MAFB acts as the global genomic anchor, while MAF acts as a localized modifier and a molecular brake on acute inflammatory chemokines, and establish that an autocrine IL-10 loop, selectively driven by MAFB, is mandatory for both the initial acquisition and the ongoing maintenance of human anti-inflammatory monocyte-derived macrophage identity.

## RESULTS

### MAF and MAFB are co-expressed and physically associate in human M-CSF-dependent M-MØ

To investigate the relative contribution of MAFB and MAF to macrophage transcriptional identity, we initially assessed their baseline expression in human monocyte-derived macrophages generated in the presence of M-CSF (anti-inflammatory M-MØ) or GM-CSF (pro-inflammatory GM-MØ). Analysis of independent transcriptomic datasets ^48^ revealed that *MAFB* and *MAF* are highly and preferentially expressed in anti-inflammatory M-MØ compared to GM-MØ, with *MAFB* consistently displaying higher read counts (Figure 1A). This expression pattern was confirmed at the protein level, as M-MØ express significantly elevated levels of both MAFB and MAF proteins (Figure 1B).

**Figure 1.**
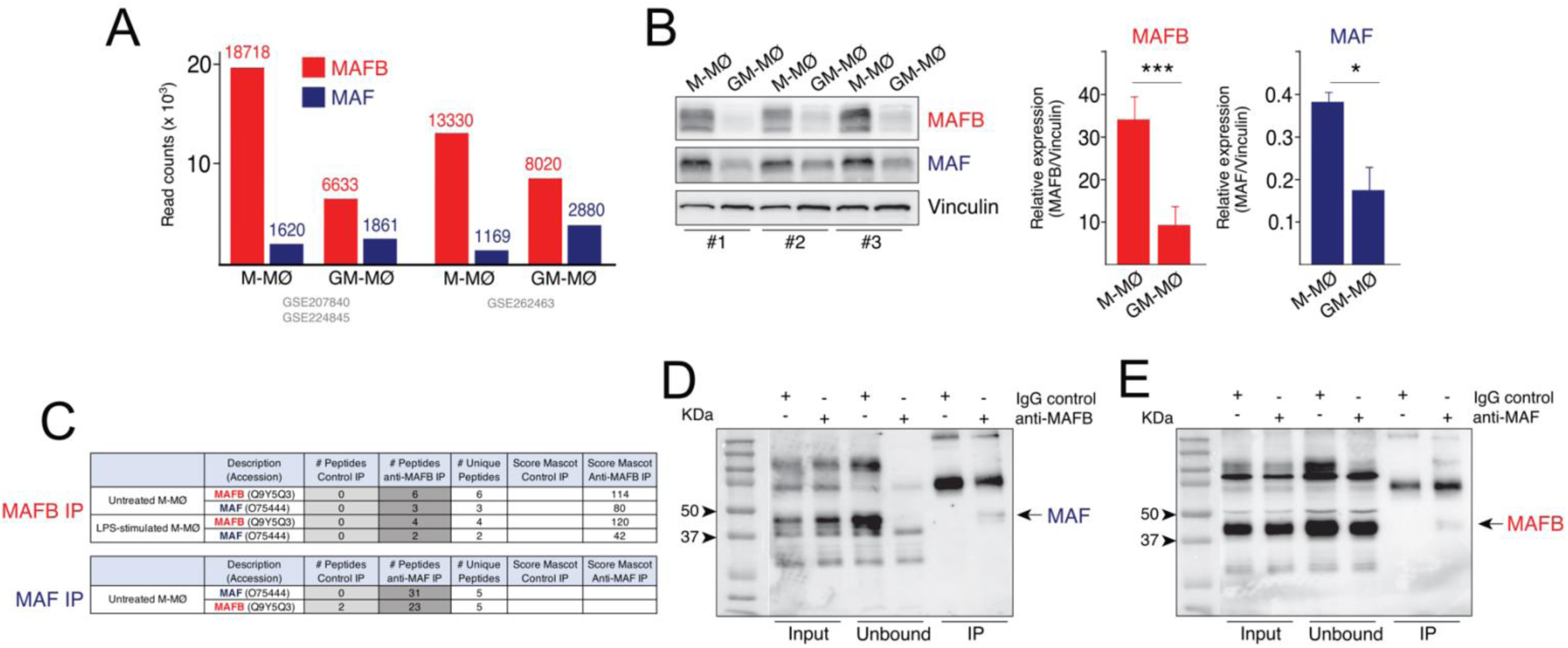
**Expression and physical association of MAFB and MAF in human M-MØ**. **A**. Relative mRNA expression (read counts) of MAFB (red) and MAF (blue) in human M-MØ and GM-MØ across independent GEO datasets (GSE207840/GSE224845 and GSE262463). **B**. MAFB and MAF protein expression in three independent preparations of M-MØ and GM-MØ (#1-#3), as determined by Western blot (*Left panel*). Quantification relative to Vinculin (*Right panels*) is shown as mean ± SEM (*, p < 0.05; ***, p < 0.001). **C**. Mass spectrometry and proteomic score metrics for anti-MAFB and anti-MAF co-immunoprecipitation experiments in human M-MØ (ProteomeXchange Consortium, PRIDE PXD083076). **D-E**. Validation of the physical interaction by endogenous co-immunoprecipitation from M-MØ lysates. Western blots show Input, Unbound, and IP fractions using anti-MAFB (**D**) or anti-MAF (**E**) antibodies, alongside control IgG.

To evaluate whether MAFB and MAF physically associate, we performed proteomic profiling of endogenous MAFB-containing complexes isolated from resting and LPS-stimulated M-MØ. Mass spectrometry analysis (PRIDE repository identifier PXD083076) identified MAF as a significant co-enriching partner under both homeostatic and inflammatory conditions (Figure 1C). Reciprocal proteomic profiling of endogenous MAF immunoprecipitates confirmed the presence of MAFB-derived peptides, albeit with a lower relative spectral count (Figure 1C). Endogenous co-immunoprecipitation (co-IP) followed by Western blotting validated this physical interaction, as MAFB immunoprecipitation pulled down MAF protein (Figure 1D) and, reciprocally, MAF pull-down enriched for MAFB protein (Figure 1E). Notably, immunoprecipitation of MAFB completely depleted MAF from the unbound supernatant, whereas a pool of MAFB remained after MAF clearance (Figure 1D-E), likely reflecting the higher baseline stoichiometry of MAFB protein in human M-MØ. Thus, MAFB and MAF form a stable, physical heterodimeric complex in human M-CSF-dependent anti-inflammatory monocyte-derived macrophages.

### MAF and MAFB distinctly orchestrate the global and identity-defining anti-inflammatory transcriptome of M-MØ

To define the specific and overlapping gene networks governed by each factor, we performed targeted siRNA-mediated silencing of *MAFB* (ΔMAFB-M-MØ) or *MAF* (ΔMAF-M-MØ) ^37^ in fully differentiated M-MØ for 24 and 36 hours, followed by transcriptomic profiling (Figure 2A-B; Supplementary Figure 1A). Global differential expression analysis (DEG) identified co-regulated target modules (genes MAFB/MAF-dependent or MAFB/MAF-repressed), factor-specific target gene sets (dependent or repressed by either MAFB or MAF), and a discrete subset of oppositely regulated genes (Figure 2C-D; Supplementary Figure 1B). Functional annotation via Enrichr ^49^ highlighted that MAFB-dependent genes were strongly associated with *MERTK* and *CSF1R* co-expression, IL-10 signaling pathways, and pro-tumoral macrophage signatures ^50^, whereas MAF-repressed genes were specifically linked to pro-inflammatory TNF and IFN signaling cascades (Supplementary Figure 1C). Thus, although MAFB and MAF share a significant portion of gene targets, they maintain specialized, non-redundant transcriptional tasks in human M-MØ.

**Figure 2.**
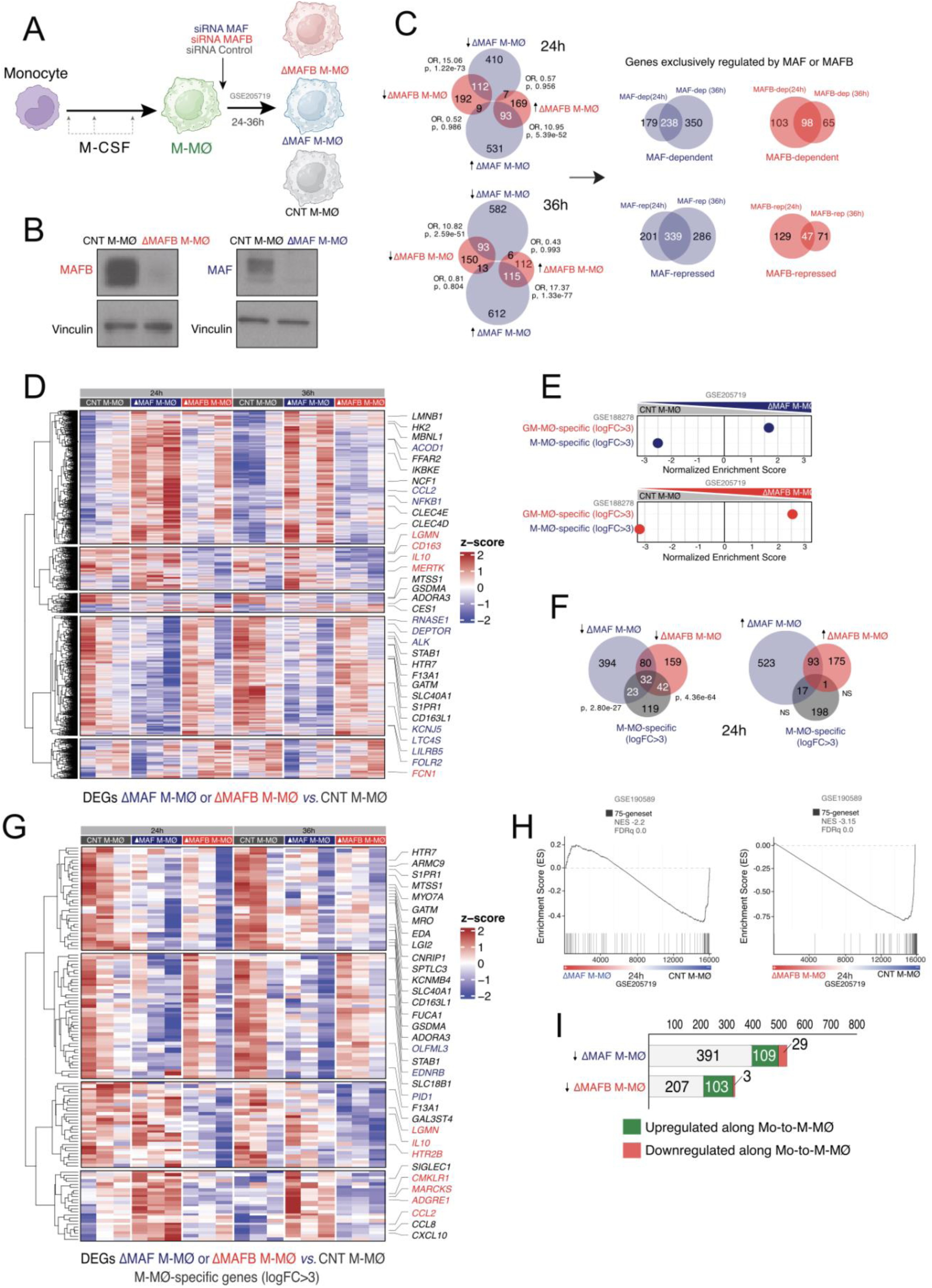
**Transcriptomic profiling of MAFB- and MAF-dependent gene networks in human M-MØ**. **A**. Schematic of the experimental workflow for siRNA-mediated silencing of MAF (ΔMAF M-MØ) or MAFB (ΔMAFB M-MØ) in M-MØ over 24−36 h. **B**. MAFB and MAF levels in ΔMAFB M-MØ, ΔMAF M-MØ and CNT M-MØ, as determined by Western blot. **C**. Venn diagrams (Hypergeometric test, with Odds Ratio, OR, and p-value) indicating overlapping, unique, and oppositely regulated differentially expressed genes (DEG) at 24 h and 36 h post-silencing in ΔMAF M-MØ and ΔMAFB M-MØ. **D**. Supervised hierarchical clustering heatmap of all identified DEG across conditions, highlighting specific genes. **E-F**. GSEA normalized enrichment scores (**E**) and Venn diagram overlaps (Hypergeometric test) (**F**) showing the selective loss of the M-MØ identity signature (M-MØ-specific, logFC>3), and enrichment of GM-MØ-specific transcripts, in ΔMAF M-MØ and ΔMAFB M-MØ. In GSEA, solid colors represent FDRq=0. **G**. Heatmap focused on the expression kinetics of core M-MØ-specific genes following knockdown of each factor. **H**. GSEA showing significant negative enrichment of MAFB-bound gene sets within the ΔMAF M-MØ and ΔMAFB M-MØ transcriptomic profiles. Normalized enrichment score and FDRq values are indicated. **I**. Distribution of down-and up-regulated DEG in ΔMAF M-MØ and ΔMAFB M-MØ relative to their physiological expression trajectory during monocyte-to-M-MØ differentiation.

Gene Set Enrichment Analysis (GSEA) ^51^ showed that silencing either factor significantly compromised the core M-MØ-specific transcriptome and shifted the global profile toward a pro-inflammatory GM-MØ-like signature (Figure 2E; Supplementary Figure 1D; Supplementary Table S1), a result supported after overlap analysis with a reference core M-MØ-specific gene set (logFC>3, representing transcripts at least 8-fold higher in M-MØ than in GM-MØ) ^48^ (Figure 2F-G; Supplementary Figure 1E-F). At 24 hours post-transfection, MAF depletion downregulated 25% of the core M-MØ identity signature ^37^^52^, while MAFB knockdown impaired the expression of 34% of these genes (Figure 2F-G). Furthermore, deficiency in either factor blunted the expression of a curated 75-gene set of direct MAFB targets ^37^ (75-geneset, Figure 2H), and genes pathologically upregulated in M-MØ from patients with Multicentric Carpotarsal Osteolysis (MCTO), a disease driven by MAFB-stabilizing mutations ^53−55^ (Supplementary Figure 1G). Furthermore, the majority of transcripts downregulated in ΔMAFB-M-MØ or ΔMAF-M-MØ corresponded to genes exclusively upregulated during monocyte-to-M-MØ differentiation (Figure 2I). Collectively, these results reveal that MAFB and MAF jointly cooperate to drive the acquisition and maintenance of the mature M-MØ anti-inflammatory transcriptome while actively repress the pro-inflammatory GM-MØ signature.

### Genome-wide profiling identifies MAFB as the primary genomic anchor of M-MØ identity

To map the elements that underpin the transcriptomic shifts in the silencing experiments, we conducted genome-wide Cleavage Under Targets and Tagmentation (CUT&Tag) profiling using anti-MAFB and anti-MAF antibodies. MAFB occupied a broad genomic landscape encompassing 4843 unique gene-assigned peaks, whereas MAF binding was restricted to a smaller, specific subset of targets (Figure 3A-C; Supplementary Tables S2-S3). Both factors localize predominantly within promoter/transcription start site (TSS) regions (44.0% for MAFB, 50.7% for MAF) and intronic regions (33.7% for MAFB, 23.8% for MAF) (Figure 3A). *De novo* motif analysis identified a significant enrichment of Maf Recognition Elements (MARE), AP-1 binding motifs, and cooperative elements like Sp1 and NF-Y motifs within the identified peaks (Figure 3B).

**Figure 3.**
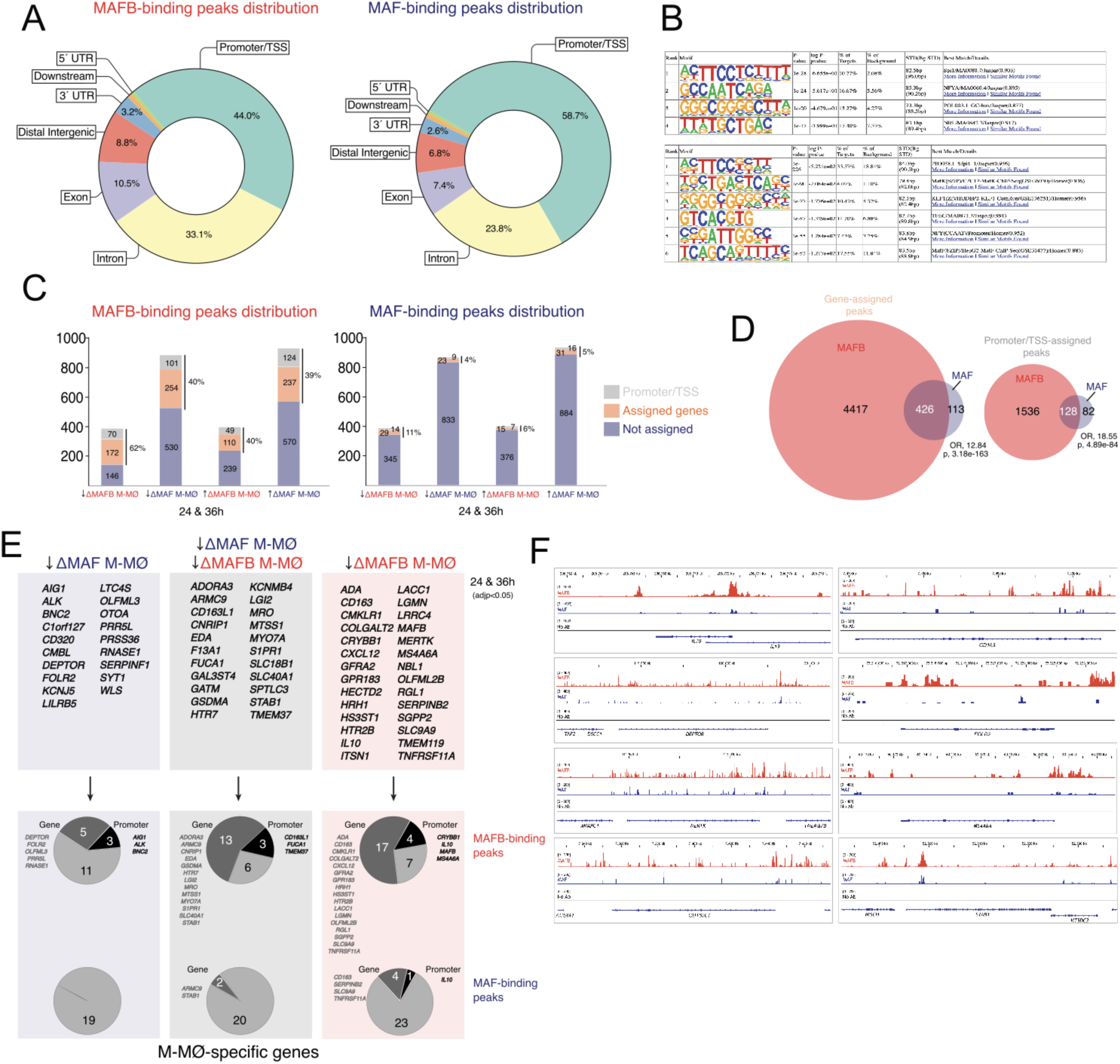
**Genome-wide CUT&Tag profiling and direct target definition for MAFB and MAF in human M-MØ**. **A**. Genomic feature distribution (promoter/TSS, intron, exon, intergenic, UTR) of MAFB- and MAF-binding peaks. **B**. High-scoring transcription factor binding motif matrices enriched within MAFB and MAF peak sites, as determined by Homer analysis. **C**. Absolute numbers and proportion of silencing-responsive DEG directly assigned to a physical binding peak or promoter/TSS element. **D**. Venn diagrams (Hypergeometric test, with OR and p-value) showing the overlap of all gene-assigned peaks and promoter/TSS sites between MAFB and MAF. **E**. Correlation analysis of M-MØ-specific identity genes downregulated upon factor silencing against physical MAFB and MAF peak/promoter occupancy, with lists highlighting specific targets. **F**. Representative IGV genome browser tracks showing raw CUT&Tag peak profiles for MAFB (red) and MAF (blue) across key macrophage loci (*IL10, CD163, DEPTOR, FOLR2, MERTK, MS4A4A, CD163L1, STAB1*), with No-Antibody controls shown in gray.

To distinguish direct transcriptional control from indirect downstream effects, we integrated genome-wide occupancy with DEG lists from ΔMAFB-M-MØ and ΔMAF-M-MØ, and found that MAFB directly binds a significant proportion of genes downregulated by either MAFB or MAF silencing, whereas independent MAF-binding peaks were markedly less frequent across the same gene cohorts (Figure 3C). Specifically, 62% of genes downregulated in ΔMAFB-M-MØ were directly assigned to a physical MAFB peak, with 20% mapping explicitly to the proximal promoter/TSS region (Figure 3C). Moreover, 40% of the transcripts upregulated in ΔMAFB-M-MØ harbored MAFB-binding sites, with 12% mapping to proximal regulatory regions (Figure 3C). While these percentages were considerably lower for independent MAF-binding peaks, 40% of the transcripts modulated upon MAF silencing also contained MAFB-binding elements (Figure 3C). This considerable structural overlap implies that a substantial proportion of MAF-regulated genes exhibit concurrent physical occupancy by MAFB, reinforcing the physiological relevance of their physical heterodimerization. In fact, the two factors shared a significant core of 426 overlapping gene peaks (*Odds Ratio*=12.84, *p*=3.18e-163) and 128 shared promoter/TSS assignments (*Odds Ratio*=18.55, *p*=4.89e-84) (Figure 3D), and core M-MØ identity could be categorized into three classes: 1) Direct shared targets (genes downregulated in ΔMAFB-M-MØ and ΔMAF-M-MØ and concurrently bound by both MAFB and MAF at their promoter and distal loci, including *STAB1*, *F13A1*, *S1PR1* and *CADM1*); 2) Direct MAFB-specific targets (downregulated exclusively in ΔMAFB-M-MØ and physically occupied solely by MAFB, like *CD163*, *MS4A4A*, *MERTK* and *IL10*); and 3) Direct MAF-specific targets (only downregulated in ΔMAF-M-MØ and with selective MAF promoter or gene binding, including *FOLR2*, *DEPTOR* and *LILRB5*).

These divergent regulatory patterns were maintained at 24 and 36h post-silencing (Figure 3E) and were largely validated in independent biological samples (Supplementary Figure 2A). Analysis of human intestinal macrophage subsets ^56^ provided strong *in vivo* translation of these findings, as *CD163* and *IL10* expression levels correlated with *MAFB* expression, while *FOLR2* levels mostly mirrored *MAF* expression (Supplementary Figure 2B). Tracking visualization of representative gene loci explicitly illustrated these binding types (Figure 3F). Robust MAFB-binding was prominent across the *IL10*, *CD163*, *MERTK*, and *MS4A4A* loci. Interestingly, while *IL10* and *CD163* exhibited distinct physical peak signals for both MAFB and MAF, their functional expression was entirely dependent on the presence of MAFB, demonstrating a clear functional asymmetry within shared genomic regions. Conversely, the loci for the M-MØ-specific *FOLR2* and *DEPTOR* genes, whose expression is primarily MAF-dependent, also demonstrated peak enrichment for MAF and MAFB.

### Shared and opposing expression of MAFB-and MAF-dependent genes *in vivo* and across tissue macrophage niches

We next intersected the MAFB and MAF regulons with published bulk and single-cell transcriptional signatures from tissue-resident macrophages (TRM) across homeostatic and pathological settings. The MAFB-dependent gene set was over-represented in “long-term” TRM (MoMac-VERSE Cluster#2) [60], in TLF^+^-like heart and liver TRM subsets ^57^, and anti-inflammatory gut M2 macrophages ^56^ (Figure 4A; Supplementary Figure 3A). Conversely, the MAF-dependent gene set was uniquely enriched in MoMac-VERSE Cluster#8 ^58^, while MAF-repressed genes were enriched in pro-inflammatory gut M1 macrophages ^56^ (Figure 4A). In the lung, the MAFB-dependent program associated with Interstitial FOLR2^+^ and SPP1^+^ macrophage subsets ^59^, whereas mature Alveolar macrophages were specifically enriched for MAFB-repressed genes ^59^ (Figure 4A). In tumoral datasets, the MAFB-dependent regulon aligned with pro-tumoral “Large TAM” from colorectal liver metastases (TAM CRC-LM) ^50^, whereas the “Small TAM” signature was enriched for MAF-dependent and MAFB-repressed gene modules (Figure 4A). Likewise, in severe COVID-19, the MAFB-dependent program mapped precisely to profibrotic monocyte-derived lung macrophage subsets (CD163^+^/LGMN^+^, MoAM3) ^60,61^ whereas the MAF-dependent module was restricted to Alveolar macrophages from infected lungs (Figure 4A). Furthermore, early inflammatory infiltrating monocyte-derived macrophages (MoAM1, MoAM2) were distinctly characterized by an enrichment in MAF-repressed genes (Figure 4A). The reciprocal representation of MAFB and MAF signatures was consistently confirmed across multiple external databases, including TRM/TAM modules ^62^, peritoneal resident macrophage modules ^63^, murine TRM profiles ^64^ (Supplementary Figure 3B-E), and TAM datasets from various malignancies ^50,65,66^ (Supplementary Figure 3F-I). These extensive tissue mappings, further supported by direct GSEA of the transcriptomes of ΔMAF-M-MØ versus ΔMAFB-M-MØ (Figure 4B; Supplementary Figure 3F-H), demonstrated that the shared and non-redundant modules of MAFB and MAF shape distinct homeostatic and pathological anti-inflammatory macrophage identities *in vivo*.

**Figure 4.**
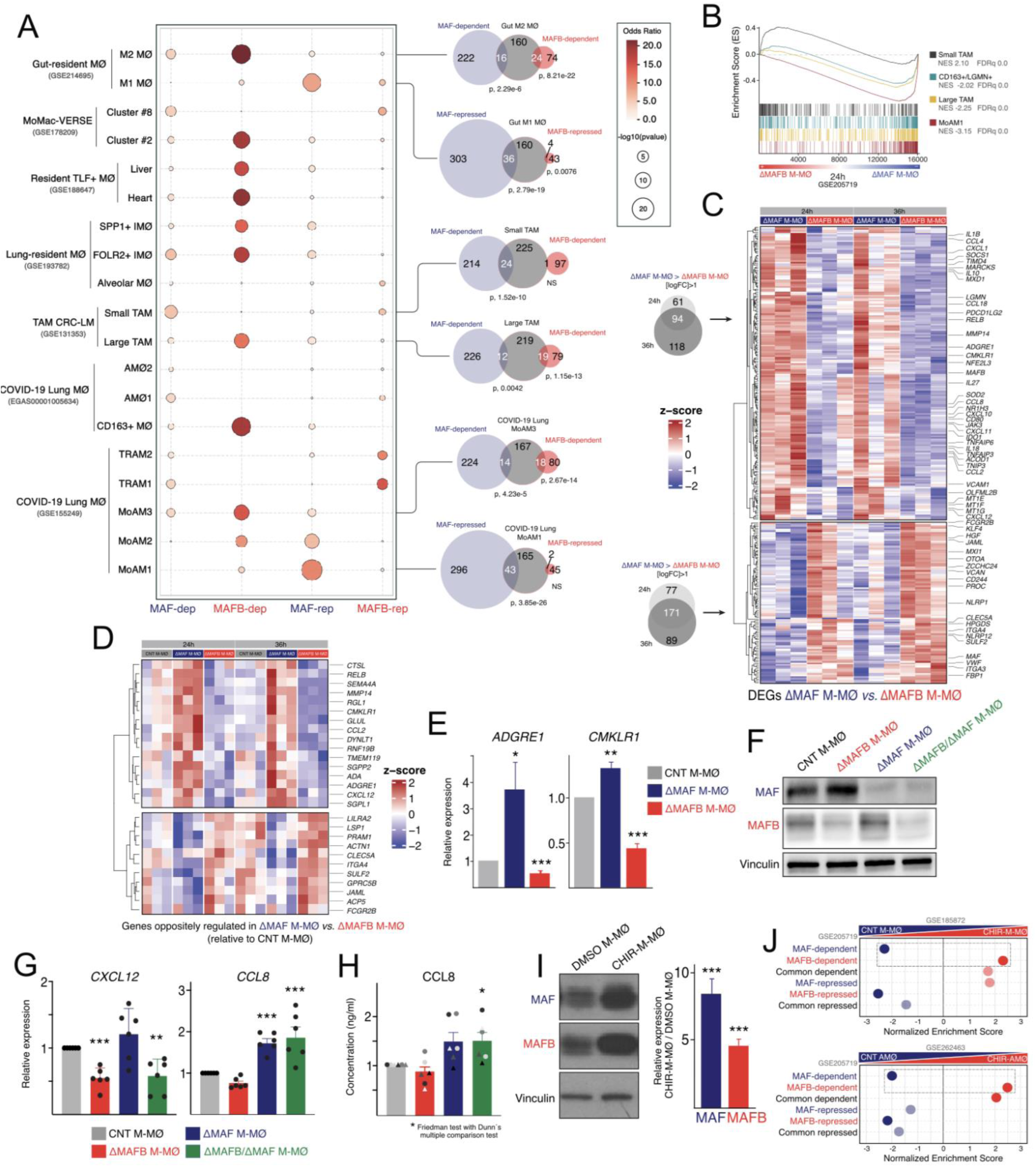
Transcriptomic divergence and differential GSK3 sensitivity of the MAFB and MAF regulatory axes. **A**. Quantitative Enrichment Analysis of MAF and MAFB Regulons. Dot plot illustrating the statistical over-representation (Hypergeometric test, with Odds Ratio, OR, and p-value) of MAFB-dependent, MAF-dependent, MAFB-repressed and MAF-repressed gene sets within the indicated macrophage signatures. The analysis was performed using a two-tailed Fisher’s Exact Test against a background of n=12000 genes. Dot size represents the statistical significance, expressed as −log_10_ (p-value). Larger dots indicate higher confidence in the association. Color Intensity represents the Odds Ratio (OR), indicating the strength of enrichment. Darker red shades denote a higher likelihood of gene set overlap compared to random expectation (max OR = 36.2). **B**. GSEA of the indicated gene sets (“Large TAM” and “Small TAM” from GSE131353, “MoAM1” from GSE155249, “CD163+/LGMN+” from EGAS00001005634) on the ranked comparison of the ΔMAF M-MØ *vs*. ΔMAFB M-MØ transcriptomes. NES and FDRq values are indicated in each case. **C**. Venn diagrams and supervised heatmap highlighting genes differentially expressed between ΔMAF M-MØ and ΔMAFB M-MØ at 24 h and 36 h post-silencing. **D** Heatmap focusing on the subset of genes showing strict opposite regulation in ΔMAF M-MØ and ΔMAFB M-MØ relative to CNT M-MØ. **E**. Relative expression of *ADGRE1* and *CMKLR1* under single-knockdown conditions, as determined by qRT-PCR on three independent samples (*, p<0.05; ***, p<0.001). **F**. Expression of MAF and MAFB protein in M-MØ after single and combined double silencing. Shown is a representative Western blot out of six independent experiments performed. **G.** Relative expression of *CXCL12* and *CCL8* under single and combined factor silencing conditions, as determined by qRT-PCR. Mean ± SEM of six independent experiments is shown (**, p<0.005; ***, p<0.001). **H**. Secretion of CCL8 (pg/ml) from ΔMAF M-MØ, ΔMAFB M-MØ, ΔMAF/ΔMAFB M-MØ and CNT M-MØ, as determined by ELISA. Mean ± SEM of six independent macrophage samples is shown (Friedman test with Dunńs multiple comparison test: *, p<0.05). **I**. Upregulation of MAF and MAFB protein levels in M-MØs upon CHIR99021-mediated GSK3 inhibition, as determined by Western blot. Mean ± SEM of three independent experiments is shown (***, p<0.001). **J.** GSEA normalized enrichment scores illustrating the opposite response profiles of MAF-dependent/repressed and MAFB-dependent/repressed gene sets in human M-MØs (*upper panel*) and alveolar macrophages (*lower panel*) upon GSK3 pathway inhibition.

### Upstream GSK3 signaling tunes the opposing transcriptional outputs of MAFB and MAF

Direct comparison between ΔMAFB-M-MØ and ΔMAF-M-MØ transcriptomes revealed the existence of genes oppositely regulated by both factors, including genes coding for homeostatic components (*IL10*, *ADGRE1*, *CMKLR1*) and key monocyte-(*CCL2, CCL8*) or neutrophil-recruiting (*CXCL1*) chemokines, whose opposite regulation by MAFB and MAF was validated in independent samples for (Figure 4C-E). To evaluate potential functional redundancies or hierarchies, both factors were simultaneously silenced in M-MØ (Figure 4F; Supplementary Figure 4A-B) and revealed gene-specific epistatic patterns: MAFB was predominat in driving *CXCL12* expression, while MAF was a non-redundant repressor of *CCL8* expression and CCL8 chemokine secretion (Figure 4G-H). Combined, these results imply that MAF and MAFB cooperate on a large portion of the genome but also exert gene-specific, and occasionally antagonistic, transcriptional actions. Furthermore, although pharmacological inhibition of Glycogen Synthase Kinase 3 (GSK3) upregulated MAFB and MAF at the protein level ^67,68^ (Figure 4I), their respective specific gene signatures (MAF-dependent, MAFB-dependent) displayed opposite sensitivity to GSK3 inhibition in M-MØ and other myeloid cells, (Figure 4J; Supplementary Figure 4C-D). Therefore, in line with their opposite effects on a set of M-MØ identity genes, the balance between the MAF and MAFB regulons can be differentiatially tuned by upstream GSK3 signaling.

### MAF acts as a non-redundant rheostat that suppresses the global macrophage pro-inflammatory activation program

As exposure of M-MØ to TLR ligands or viral particles rapidly downregulates MAF and MAFB expression ^69^ (Supplementary Figure 5A-C). we examined the contribution of their respective targets to macrophage activation by various inflammatory stimuli, including TLR ligands, SARS-CoV-2, TNF, *Mycobacterium tuberculosis* or IFNγ (Figure 5A, Supplementary Figure 5D). M-MØ activation resulted in a substantial increase in the expression of MAF-repressed genes, but not MAFB-repressed genes (Figure 5A, Supplementary Figure 5D), suggesting that MAF preserves the M-MØ homeostatic identity by actively restraining pro-inflammatory activation. Strikingly, MAF silencing closely mimicked M-MØ activation (Figure 5B-C), with a shared “core” of 120-180 genes commonly upregulated by LPS, CL264, and after MAF knockdown ^70,71^ (Figure 5D), including critical inflammatory nodes (*ACOD1*, *CCL5*, *CXCL1, NFKB1*, *NFKB2*, *RELB*, *TNF*, and *TNFAIP3*) (Figure 5D) and covering over 65% of the whole MAF-repressed gene set. This result identifies MAF, and not MAFB, as a molecular brake on acute inflammation. In fact, comparing these modules against classical “trained immunity” signatures ^72^ revealed that MAF silencing in M-MØ recaptures ∼50% of the transcriptional intensity of β-glucan-trained monocytes ^72^ (Scores: 2.24 at 2h; 2.19 at 4h) and hyper-inflammatory sepsis monocytes ^73,74^, features not observed with the MAFB-repressed core (0.67 at 2h, 0.65 at 4h) (Figure 5E; Supplementary Figure 5E).

**Figure 5.**
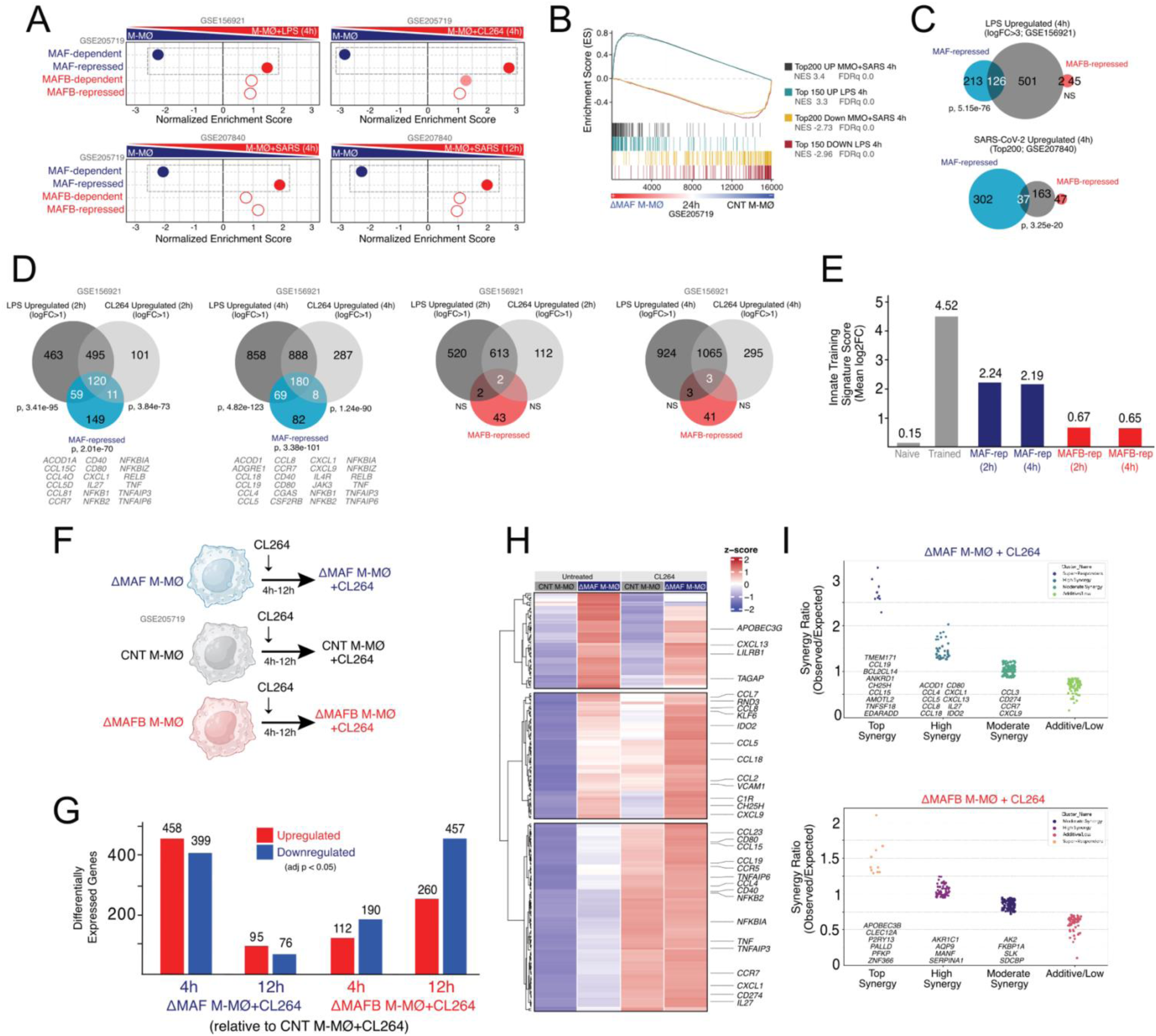
MAF silencing mimicks M-MØ activation and primes M-MØ for stronger inflammatory responses. **A.** Summary of GSEA of the indicated gene sets on the ranked comparison of the transcriptomes of M-MØ+LPS (4h) (GSE156921), M-MØ+CL264 (4h) (GSE205719), M-MØ+SARS (4h) (GSE207840) or M-MØ+SARS (12h) (GSE207840) versus the transcriptome of resting M-MØ. NES and FDRq values are indicated in each case (Solid-colored dots, FDRq=0; Light-colored dots, FDRq<0.250; Empty dots, Non significant FDRq). **B**. GSEA of the indicated gene sets (“Top150 LPS UP (4h)” and “Top150 LPS DOWN (4h)” from GSE156921, “Top200 SARS-CoV-2 UP (4h)” and “Top200 SARS-CoV-2 DOWN (4h)” from GSE207840) on the ranked comparison of the ΔMAF M-MØ *vs*. CNT M-MØ transcriptomes. NES and FDRq values are indicated in each case. **C**. Venn diagrams showing the overlap (Hypergeometric test, with p-value) of the most strongly upregulated genes in M-MØ exposed for 4 hours to either LPS (LPS Upregulated (4h), logFC>3; GSE156921) (*Upper panel*) or SARS-CoV-2 (SARS-CoV-2 Upregulated (4h), Top200; GSE207840) (*Lower panel*), with the MAF-repressed and MAFB-repressed gene sets. **D**. Venn diagrams showing the overlap (Hypergeometric distribution test against a genomic background of 12.000 genes, with indication of p-value) of the genes upregulated (adj-p<0.05; log_2_FC>1) in M-MØ exposed for 2 or 4 hours to either LPS or CL264 (GSE156921) with the MAF-repressed and MAFB-repressed gene sets. Selected overlapping genes are indicated in the case of the MAF-repressed gene set. **E**. Quantitative validation of the “pseudo-trained” transcriptional state in ΔMAF M-MØ and ΔMAFB M-MØ. The Innate Training Signature Score represents the Mean log_2_Fold Change (Mean log_2_FC) of the exclusive regulatory modules for MAF and MAFB that overlap with the core TLR4/7-inducible program. To provide a physiological baseline, the reference scores for “Naive” and “Trained” states (grey-shaded bars) represent the average induction of training-associated genes in β-glucan-stimulated monocytes ^72,74^. Values above bars indicate the Innate Training Signature Score for the indicated conditions, and represent the mean induction of the triple-overlap gene cores (Agonist 2h/4h & MAF/MAFB-repressed). **F**. Schematic of the experimental workflow for CL264 activation of ΔMAF M-MØ, ΔMAFB M-MØ or CNT M-MØ over 4−12 hours. **G**. Number of Differentially Expressed Genes (DEG, adjp<0.05) between the transcriptomes of CL264-treated ΔMAF M-MØ (ΔMAF M-MØ+CL264) or ΔMAFB M-MØ+CL264 (ΔMAFB M-MØ+CL264) and CL264-treated CNT M-MØ (CNT M-MØ+CL264). **H**. Heatmap of the expression of genes exhibiting an additive or synergistic response to MAF knockdown and exposure to CL264. Selected genes are indicated. **I**. Identification of genes exhibiting various degrees of synergy upon exposure to CL264 and knockdown of MAF (*Upper panel*) or MAFB (*Lower panel*). Shown is the Synergy Ratio, that represent the ratio between the Observed expression of each gene in ΔMAF M-MØ+CL264 and its Expected expression (additive) according to its levels in ΔMAF M-MØ and in CNT M-MØ+CL264.

To functionally test the ability of MAF to affect the intensity of the macrophage proinflammatory response, we analyzed the effect of MAF (ΔMAF-M-MØ) or MAFB (ΔMAFB-M-MØ) silencing on the M-MØ response to the TLR7 ligand CL264 for 4-12 hours (Figure 5F). While the total number of responsive DEG was similar across groups (Supplementary Figure 5F), the transcriptomes of CL264-treated ΔMAF-M-MØ and ΔMAFB-M-MØ exhibited qualitative differences from CL264-treated control cells (Figure 5G). Crucially, MAF silencing triggered a profound synergistic response with TLR7 stimulation for over 80% of the MAF-repressed regulon (Figure 5H-I), whereas a much weaker synergy was seen in the case of MAFB silencing (Figure 5H-I; Supplementary Figure 5G). Therefore, MAF and MAFB constitute non-redundant checkpoints that preserve M-MØ homeostasis.

### Transcriptional divergence dictates distinct macrophage effector functions

We next assessed whether the distinct transcriptional programs of MAF and MAFB translate into specific M-MØ effector functions, and functional assays confirmed the operational segregation between the two factors. MAF deletion increased basal and TLR-induced *CCL2* gene expression and CCL2 protein release, as well as TLR-induced *CCL8* expression and secretion (Figure 6A-B). Further, MAF and MAFB deletion had opposite effects on the CL264-induced expression of *IL1A* and *IL23A* (Figure 6A-B). In homeostatic clearance assays, MAFB was required for apoptotic cell efferocytosis, matching the global decrease in efferocytosis-related transcripts in ΔMAFB-M-MØ (Figure 6C; Supplementary Figure 6A), whereas phagocytosis of bacterial particles solely relied onj MAF (Figure 6D). Finally, MLR assay showed that MAFB deficiency, but not MAF deficiency, affected the capacity of macrophages to induce allogeneic CD4+ T cell proliferation (Figure 6E; Supplementary Figure 6B). However, both MAF or MAFB silencing altered the cytokine signature of responding T cells, decreasing IL-10 production while increasing IFN-ɶ production (Figure 6F-G). Therefore, despite their shared capacity to shape the anti-inflammatory profile, MAF and MAFB exhibit transcriptomic differences and exert non-redundant roles in human monocyte-derived macrophages.

**Figure 6.**
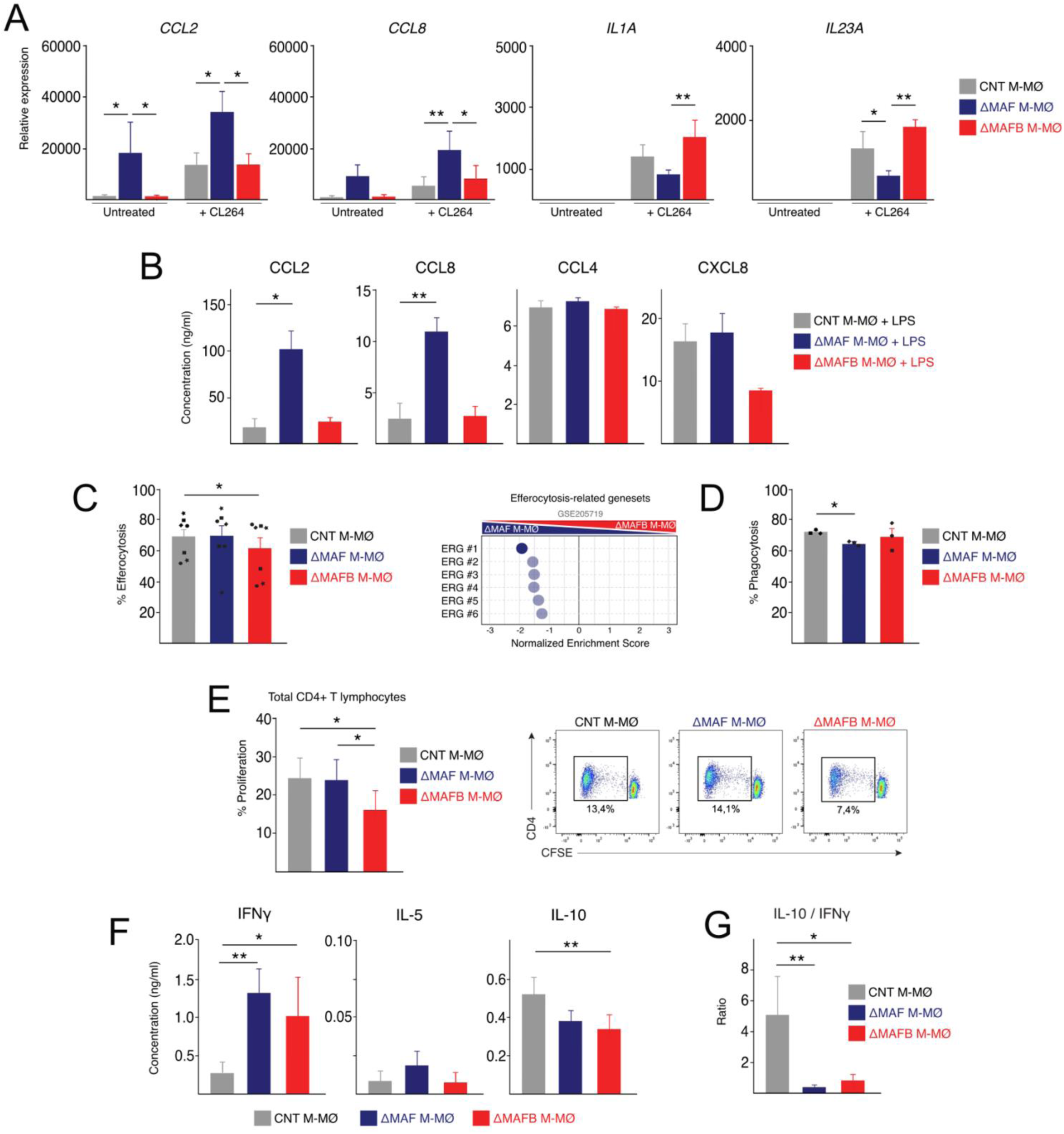
Distinct functional consequences of MAF and MAFB knock-down. **A.** Relative mRNA expression of the indicated genes in CNT M-MØ, ΔMAF M-MØ, and ΔMAFB M-MØ before and after exposure to CL264, as determined by RT-PCR. Mean ± SEM of three independent samples are shown (Paired Student’s t-test: *, p<0.05; **, p<0.01). **B**. Production of the indicated cytokines by CNT M-MØ, ΔMAF M-MØ, and ΔMAFB M-MØ after exposure to LPS, as determined by ELISA. Mean ± SEM of three independent samples are shown (paired Student’s t-test: *, p<0.05; **, p<0.01). **C**. Efferocytosis capacity of CNT M-MØ, ΔMAF M-MØ and ΔMAFB M-MØ, as determined by flow cytometry using staurosporine-induced CellTrace Violet-labeled apoptotic Jurkat cells. The percentage of positive cells is shown (*Left panel*). Mean ± SEM of seven independent samples are shown (one-way ANOVA Repeated Measures with Dunnett’s multiple comparisons test: *, p<0.05). The *Right panel* shows a summary of GSEA of various independent Efferocytosis-related gene sets on the ranked comparison of the transcriptomes of ΔMAF M-MØ vs. ΔMAFB M-MØ (GSE205719). NES and FDRq values are indicated in each case (Solid-colored dots, FDRq=0; Light-colored dots, FDRq<0.250). **D**. Phagocytosis of pHRodo-labeled bacterial particles by CNT M-MØ, ΔMAF M-MØ and ΔMAFB M-MØ, as determined by flow cytometry. Mean ± SEM of three independent samples is shown (one-way ANOVA Repeated Measures with Dunnett’s multiple comparisons test: *, p<0.05). **E**. Allogeneic CD4^+^ T lymphocyte proliferation promoted by CNT M-MØ, ΔMAF M-MØ or ΔMAFB M-MØ, as determined by ^3^H-Thymidine incorporation and using three independent preparations of M-MØ (donor#1-3). For each experiment, Mean ± SEM of four replicates is shown (*, p <0.05; ***, p <0.001). **F**. Proliferation of allogeneic CD4^+^ T lymphocyte and Treg promoted by CNT M-MØ, ΔMAF M-MØ or ΔMAFB M-MØ, as determined by flow cytometry on CFSE (carboxyfluorescein succinimidyl ester)-labeled cells to track T cell division. Data are shown as mean ± SEM from five independent experiments using macrophages from three independent donors and T lymphocytes from two independent donors. Each macrophage donor was combined with T lymphocytes from one donor, and two of the macrophage donors were also combined with T lymphocytes from the second donor, resulting in five independent donor combinations. Statistical analysis done with paired Student’s t-test (*p < 0.05) for the indicated comparisons.

### The MAFB/MAF complex establishes an autocrine IL-10 loop required for acquisition and maintenance of M-MØ identity

M-MØ are classically defined as “non-polarized” monocyte-derived macrophages *in vitro*, yet their transcriptional signature resembles TRM ^67^, tumor-associated macrophages and pro-fibrotic lung macrophages from severe COVID-19 ^69,75^. Given *IL10* expression in M-MØ is predominantly driven by MAFB ^22^ (Figure 2G; Supplementary Figure 2A), and that both MAFB and MAF physically occupy the proximal regulatory region of the *IL10* locus (Figure 3F), we hypothesized that the M-MØ transcriptional identity could be shaped by endogenously produced IL-10. To test this hypothesis, human monocytes were differentiated into M-MØ either in the continuous presence of a neutralizing anti-IL-10 antibody (M-MØ+anti-IL10, to assess identity *acquisition*), or exposed to the blocking antibody only for the final 48 hours of differentiation (M-MØ+anti-IL10*, to assess identity *maintenance*), and the transcriptional profile of the resulting cells was determined. In the *maintenance* model (M-MØ+anti-IL10*, Figure 7A), the short-term blockade of autocrine IL-10 diminished *FOLR2* expression, and impaired the global expression of core M-MØ-specific genes, homeostatic MAF- and MAFB-dependent targets (Figure 7B-D; Supplementary Figure 6C), and the defining signatures of M2-like gut macrophages, long-term TRM, and Large TAMs (Supplementary Figure 6D-G).

**Figure 7.**
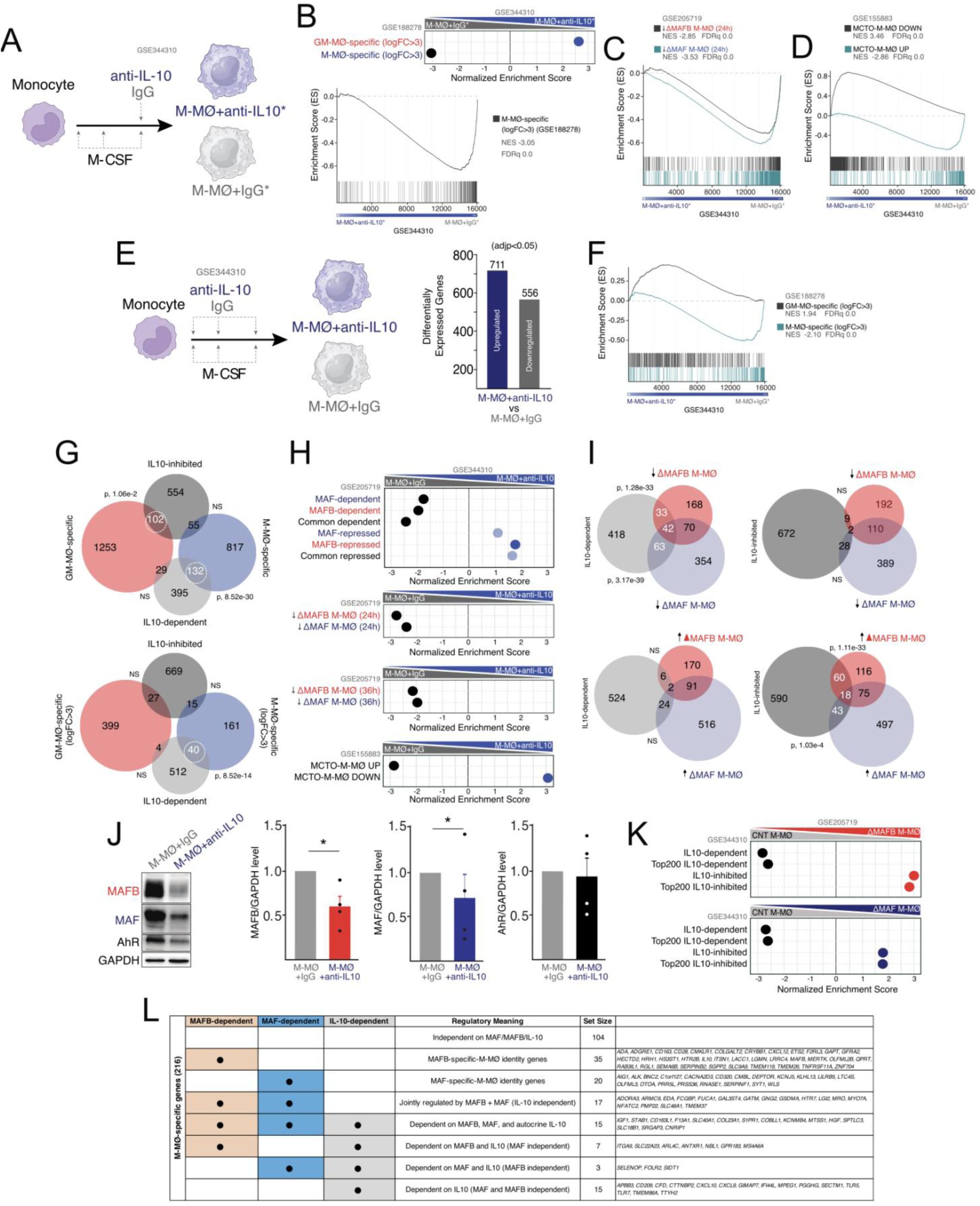
IL-10 contribution to the maintenance and acquisition of the M-MØ transcriptional identity. **A**. Schematic of the anti-IL-10 treatment (1 single dose) during the monocyte-to-M-MØ differentiation process to generate M-MØ+anti-IL10*. **B**. Summary of GSEA of GM-MØ- and M-MØ-specific gene sets (GSE188278) on the ranked comparison of the transcriptomes of M-MØ+IgG* vs. M-MØ+anti-IL10* (GSE344310). The GSEA profile for the M-MØ-specific gene set is shown below. **C**. GSEA of the genes downregulated upon MAF or MAFB knockdown (GSE205719) on the ranked comparison of the transcriptomes of M-MØ+IgG* vs. M-MØ+anti-IL10*. **D**. GSEA of the genes whose expression is altered in MCTO M-MØ (GSE155883) on the ranked comparison of the transcriptomes of M- MØ+IgG* vs. M-MØ+anti-IL10*. **E**. Schematic of the anti-IL-10 treatment (3 doses) during the monocyte-to-M-MØ differentiation process to generate M-MØ+anti-IL10. The number of DEG between M-MØ+anti-IL10 and M-MØ+IgG is indicated (GSE344310). **F**. GSEA of GM-MØ- and M-MØ-specific gene sets (GSE188278) on the ranked comparison of the transcriptomes of M-MØ+IgG vs. M-MØ+anti-IL10. **G**. Venn diagrams showing the overlap (Hypergeometric test, with p-value) of the genes upregulated (IL-10-inhibited) or downregulated (IL-10-dependent) in M-MØ+anti-IL10 (adj-p<0.05) with GM-MØ- and M-MØ-specific gene sets (GSE188278). **H**. Summary of GSEA of the indicated gene sets on the ranked comparison of the transcriptomes of M-MØ+IgG vs. M-MØ+anti-IL10. The GSEA profile for the M-MØ-specific gene set is shown below. **I**. Venn diagrams showing the overlap (Hypergeometric test, with p-value) of the genes upregulated (IL-10-inhibited) or downregulated (IL-10-dependent) in M-MØ+anti-IL10 (adj-p<0.05) with the genes downregulated or upregulated in ΔMAF M-MØ or ΔMAFB M-MØ (GSE205719). **J.** (*Left panel*) Expression of MAFB, MAF and AhR in M-MØ+IgG and M-MØ+anti-IL10, as determined by Western blot. GAPDH protein levels were determined as protein loading control. (*Right panel*) Mean ± SEM of the MAFB/GAPDH, MAF/GAPDH and AhR/GAPDH protein ratios from four independent experiments are shown (paired Student’s t-test: *, p<0.05). **K**. Summary of GSEA of IL10-dependent and IL10-inhibited gene sets on the ranked comparison of the transcriptomes of ΔMAFB M-MØ vs. CNT M-MØ and ΔMAF M-MØ vs. CNT M-MØ. NES and FDRq values are indicated in each case (Solid-colored dots, FDRq=0). **L.** Specific dependency (MAFB, MAF, IL-10) of the core genes defining M-MØ identity.

These results imply that autocrine IL-10 significantly contributes to maintaining the stability of the MAFB/MAF-dependent M-MØ transcriptome.

The role of IL-10 during the identity *acquisition* process (M-MØ+anti-IL10) was more robust, as its blockade altered the expression of 1267 genes (Figure 7E), abolishing the acquisition of the standard M-MØ-specific transcriptomic signature (Figure 7F), the homeostatic MAF- and MAFB-dependent regulons (Figure 7G-H), and the gene sets defining M2-like gut macrophages, and various TRM and TAM subsets (Supplementary Figure 6H-K). In fact, IL-10-dependent genes significantly overlapped with the transcripts downregulated upon MAFB or MAF silencing (Figure 7I). The continuous neutralization of IL-10 throughout the differentiation process resulted in a marked decrease of both MAFB and MAF proteins (Figure 7J), and even in Ser9-GSK3β levels (Supplementary Figure 6L), providing a clear mechanistic rationale for the profound collapse of M-MØ-specific, MAFB-dependent, and MAF-dependent gene signatures upon IL-10 deprivation. Reciprocally, GSEA demonstrated that MAFB silencing (ΔMAFB-M-MØ) triggered a dramatic reduction in the expression of IL-10-dependent genes, an effect that was significantly weaker in ΔMAF-M-MØ cells (Figure 7K). Because the downstream transcriptomic targets of autocrine IL-10 largely overlap with the MAFB/MAF-regulated cistrome, these results establish that endogenously produced IL-10 critically cooperates with MAFB/MAF to shape and sustain the anti-inflammatory M-MØ transcriptome. These findings align with the dominant role of MAFB in driving IL-10 cytokine production ^22^, and provide an integrated framework defining the intertwined factor and cytokine dependencies that establish the core identity of human M-CSF-dependent macrophages (Figure 7L).

## DISCUSSION

We report that the large MAF transcription factors MAFB and MAF heterodimerize under both homeostatic and inflammatory conditions, and cooperatively and distinctly govern the transcriptional identity and functional specialization of human monocyte-derived macrophages (M-MØ). While both factors are central coordinators of the macrophage anti-inflammatory state, yet single-factor silencing reveals a highly gene-dependent, non-redundant division of labor. MAFB serves as the primary global anchor of the M-MØ genome, whereas MAF provides localized, locus-specific tuning. At the functional level, this molecular divergence manifests as specialized effector programs: MAFB selectively orchestrates apoptotic cell efferocytosis, whereas MAF modulates standard phagocytosis, limits T-cell proliferative potency, and acts as a vital molecular brake suppressing acute pro-inflammatory chemokine hyper-induction. Moreover, although both factors physically occupy regulatory elements of the *IL10* locus, functional *IL10* expression is strictly dependent on MAFB. This is of crucial relevance, as neutralizing antibody assays indicate that endogenous IL-10 establishes an autocrine feedback loop that helps in the initial acquisition and long-term maintenance of the anti-inflammatory profile of M-MØ.

To our knowledge, we described for the first time the direct association in human macrophages of endogenous MAFB and MAF, whose interaction had been previously deduced from *in vitro* experiments using peptide arrays and Fluorescent Resonance Energy Transfer ^76,77^. The heterodimerization of MAFB and MAF in human M-MØ highlights a coordinated regulatory axis, while genome-wide CUT&Tag and transcriptomic data demonstrate that both factors are far from redundant. MAFB commands a wider genomic cistrome, binding thousands of unique loci, whereas MAF is recruited to a tightly restricted subset of target genes. This structural asymmetry suggests a model where the broader macrophage homeostatic program is mostly dictated by MAFB, while MAF is recruited to shared sites or unique loci to fine-tune specialized transcriptional inputs. The physiological relevance of this division is supported by the small subset of genes subjected to strict opposite regulation by these factors. The reciprocal dependency of genes like *ADGRE1* (coding for the macrophage marker F4/80) and *CMKLR1* exemplifies how these associated factors can act as an internal rheostat, shifting the phenotypic profile of the macrophage based on the relative composition and abundance of the heterodimer at specific promoter-enhancer elements. Similarly, the transcriptional divergence of both factors translates into distinct physiological outcomes: MAFB primarily drives apoptotic cell efferocytosis, matching the profound loss of efferocytic machinery observed upon its deletion, whereas standard bacterial phagocytosis mostly relies on MAF. It is tempting to speculate that this segregation might prevent macrophage vulnerability: by maintaining distinct transcription factor dependencies for efferocytosis and phagocytosis, macrophage would ensure that tissue-repair programs (efferocytosis) and anti-pathogen responses (phagocytosis) can be modulated independently in response to their microenvironment. Of note, upstream GSK3 signaling is capable of shifting this balance as pharmacological inhibition of GSK3 triggers an asymmetric upregulation of MAF over MAFB, what suggests that the MAFB/MAF complex is a tunable sensor that responds to extrinsic signaling pathways to reshape the anti-inflammatory landscape.

Importantly, our data also establishes that MAF, and not MAFB, restrains acute inflammatory output: MAF silencing under homeostatic conditions triggers a profound pro-inflammatory reprogramming that mimics classical TLR activation and trained immunity signatures (upregulation genes like *ACOD1*, *TNF*, and key components of the NF-κB cascade), and drives a synergistic hyper-induction of pro-inflammatory chemokines (*CCL15*, *CCL19*) when combined with TLR7 stimulation. These results indicate that MAF functions to limit the macrophage inflammatory response. However, since no inflammatory genes (*TNF*, *CCL5*, or *NFKB1*) are direct targets of MAF in the CUT&Tag data, MAF does not appear to be an active transcriptional repressor but an “upstream network gatekeeper” whose loss causes a systemic cellular stress that indirectly activates the NF-kB pathway. In fact, the ability of MAFB to bind MAF, as well as various member of the NFκB family (PRIDE repository, identifier PXD083076), suggests that MAF might modulate the activity of the NFκB pathway through its interaction with MAFB.

Our results have also revealed a clear asymmetry in controlling the *IL10* locus in human macrophages. While numerous studies in murine models and specific cell lines have frequently grouped MAFB and MAF as joint drivers of IL-10 expression ^30,31^, *IL10* gene expression is only selectively reduced upon MAFB depletion in M-MØ. Therefore, since both MAFB and MAF bind to the human *IL10* gene, MAF binding appears to be insufficient to sustain baseline *IL10* transcription in the absence of its partner, and may act as a stabilizing factor. Alternatively, the higher MAFB levels expressed by M-MØ might justify this asymmetrical dependency of *IL10* gene expression. Regardless of the mechanism, this differential dependency would allow monocyte-derived macrophages to preferentially link IL-10 expression to the lineage-stable levels of MAFB rather than the more activation-sensitive levels of MAF.

The MAFB-IL-10 axis we have identified fits with previous reports on the link between MAFB, its upstream regulator GSK3, and IL-10 gene expression ^78−8^^1^. This axis is of utmost importance because continuous or short-term blockade of IL-10 disrupts the M-MØ transcriptome, what reveals the existence of an autocrine loop that significantly contributes to the M-MØ gene signature. Thus, M-CSF drives the initial expression of the MAFB/MAF axis, which turns on endogenous IL-10 expression, creating a self-sustaining autocrine circuit. In fact, whereas short-term IL-10 blockade in fully mature M-MØ severely damages the stability of the M-MØ gene signature, the continuous IL-10 neutralization not only reduces the core anti-inflammatory cistrome of M-MØ but causes a shift toward a more pro-inflammatory GM-MØ-like state. Therefore, the M-MØ identity appears to require a continuous reinforcement by IL-10, that impacts directly on the gene networks that are MAFB- or MAF-dependent, implying the existence of nested regulatory feedback: MAFB/MAF are required to produce IL-10, and IL-10, in turn, relies on the collaborative network of downstream MAFB and MAF target genes to execute its primary homeostatic program. The fact that *FOLR2*, widely recognized as a major marker for human tissue macrophages ^57^, is the only gene significantly downregulated upon exposure to a single dose of blocking IL-10 strongly supports an important role for IL-10 in the acquisition of the gene expression profile of Tissue-Resident Macrophages. Indeed, the MAFB/IL-10-dependent gene modules map precisely onto long-term tissue-resident macrophage populations across multiple organs, including mature anti-inflammatory M2 macrophage subsets in the human gut, and interstitial FOLR2^+^ macrophages in the lung.

In conclusion, our study establishes that the physical association of MAFB and MAF, paired with the autocrine IL-10 feedback loop, constitutes a central molecular mechanism that defines the human anti-inflammatory M-MØ identity. This collaborative network ensures a highly stable homeostatic baseline while maintaining the exact functional plasticity required to interact with complex tissue microenvironments, thus highlighting the MAFB/MAF/IL-10 axis as an attractive therapeutic target for diseases caused by unregulated inflammatory responses.

### Limitations of the present study

We are aware that although the translational cross-referencing to *in vivo* single-cell data is excellent, our primary mechanistic discoveries (co-IP, silencing, CUT&Tag) rely on *in vitro* derived M-CSF-differentiated human monocyte-derived macrophages. However, we might also consider this an advantage rather than a limitation because human tissue-resident macrophages are highly scarce and difficult to isolate without triggering immediate activation artifacts, and the use of primary human monocyte-derived M-MØ allows the generation of the high cell yields required for high-input biochemistry (like endogenous co-IP/Mass Spectrometry) without chemical manipulation.

## METHODS

### Generation of human monocyte-derived macrophages *in vitro*

Buffy coats from anonymous healthy blood donors were provided by Comunidad de Madrid blood Bank. Ethical approvals for all blood processes used in this study were approved by the Centro de Investigaciones Biológicas Ethics Committee, and all experiments were carried out in accordance with the approved guidelines and regulations. Human Peripheral Blood Mononuclear Cells (PBMCs) were isolated from buffy coats over a Lymphoprep (Nycomed Pharma) gradient according to standard procedures. Monocytes were purified from PBMC by magnetic cell sorting using anti-CD14 microbeads (Miltenyi Biotec). Monocytes (>95% CD14+ cells) were cultured at 0.5 x 106 cells/ml in Roswell Park Memorial Institute (RPMI 1640, Gibco) medium supplemented with 10% fetal bovine serum (FBS, Biowest) (complete medium) for 7 days in the presence of 10 ng/ml M-CSF or 1000 U/ml GM-CSF (ImmunoTools) to generate M-CSF-polarized macrophages (M-MØ) or GM-CSF-polarized macrophages (GM-MØ) or respectively ^15,16^. Cytokines were added every two days and cells were maintained at 37°C in a humidified atmosphere with 5% CO_2_ and 21% O_2_. When indicated, macrophages were exposed to the GSK3 inhibitor CHIR99021 (10 μM), using DMSO as control, or exposed to 1-3 doses of an anti-IL-10 blocking antibody (2.5 μg/ml, R&D Systems #MAB2171-500) or an isotype-matched control antibody (IgG2b, 2.5 μg/ml, R&D Systems #MAB004). For macrophage activation, cells were treated with either 100 ng/ml CL264 (InvivoGen) or 10 ng/ml E. coli 0111:B4 lipopolysaccharide (LPS-EB Ultrapure, Invivogen). Human cytokine production was measured in culture supernatants using commercial ELISA [CCL2 (BD Biosciences, #555179), CCL4 (R&D Systems, #DY271-05), CCL8 (R&D Systems, #DY281) and CXCL8 (R&D Systems, #DY208)] and following the procedures supplied by the manufacturers.

### Quantitative PCR

Total RNA was extracted using the total RNA and protein isolation kit (Macherey-Nagel). RNA samples were reverse transcribed with High-Capacity cDNA Reverse Transcription reagents kit (Applied Biosystems) according to the manufacturer’s protocol. Quantitative PCR (qPCR) was performed with Light-Cycler 480 Probes Master (Roche Life Sciences) and TaqMan probes on a standard plate in a Light Cycler 480 instrument (Roche Diagnostics). Gene-specific oligonucleotides were designed using the Universal ProbeLibrary software (Roche Life Sciences). Results were normalized to the expression level of the endogenous references genes *TBP* and *HPRT1* and quantified using the ΔΔCT (cycle threshold) method.

### siRNA transfection

M-MØ (1 × 10^6^ cells) were transfected with a human *MAF*-specific (siRNA MAF, 50 nM; Dharmacon) and/or *MAFB*-specific siRNA (siRNA MAFB, 50 nM; Dharmacon) using HiPerFect (Qiagen). Silencer Select Negative Control No. 2 (siRNA Control, 50 nM; Dharmacon) was used as negative control siRNA. Six hours after transfection, cells were either allowed to recover from transfection in complete medium, collected 24/36 hours (24h, 36h) after transfection and processed for further analysis. Knockdown of MAF/MAFB was confirmed by Western blot.

### Co-immunoprecipitation

For identification of MAFB- or MAF-associated proteins, 6-10×10^7^ M-MØ were used per immunoprecipitation condition. Cells were lysed in cold using 50 mM Tris pH 8.0, 1 mM EDTA and 150 mM NaCl (Basal buffer), containing 1% Triton X-100, cOmplete™ Protease Inhibitor Cocktail (Merck, #11697498001) and PhosSTOP™ phosphatase inhibitor (Merck, # 4906845001) (Lysis Buffer). Benzonase (100U; Merck, # E1014-5KU) was used for removal of nucleic acid for 6h at 4°C. After centrifugation at *7000g* for 15 min and preclearing, cell lysates were incubated overnight at 4°C with 15μl of MagnaChIP A+G magnetic beads (Merck, #16-663) previously coated with the corresponding antibodies (MAFB: Rabbit polyclonal #HPA005653, Merck; MAF: E2Q5D Rabbit mAb #26745, Cell Signaling Technology) following the manufactureŕs recommendations. After extensive washing using Lysis buffer diluted 1:10 in Basal buffer, beads were resuspended in Basal buffer before proteomics analysis. Alternatively, in the case of MAFB immunoprecipitation from resting M-MØ, beads were incubated with 200 µL Glycine 0.2M pH 2.3, and the supernatant processed for proteomics analysis. Immunoprecipitation was validated by Western blot before further analysis.

### Proteomics

For proteomics analysis of the MAFB immunoprecipitates from LPS-treated M-MØ, and MAF immunoprecipitates from resting M-MØ, digestion was carried out using the Preomics iST kit following the manufactureŕs instructions. In brief, magnetic beads were treated with 50 µl of LYSE buffer for 10 minutes at 95°C, transferred to the columns, and 50 µl of DIGEST solution was added. After 2 h at 37°C, 100 µl of STOP buffer was added, and the mixture shaken and centrifuged for 2 min at *3800g*. The digested sample was washed with 100 µl of WASH1 buffer, then with 100 µl of WASH2 buffer, with 2 min centrifugation at *3800g* between washes, and eluted twice with 100 µl of ELUTE. The sample was dried in a SpeedVac and reconstituted in 20 µl of LC-LOAD. For Data-dependent acquisition (DDA) (shotgun proteomics) analysis, approximately 10 µl of the peptide digest was loaded and concentrated onto a tip (Evotip) following manufactureŕs instructions. After that, peptides were separated by nano-liquid chromatography (EvoSep One, EVOSEP) in a 15 cm chromatographic column (PepSep C18 15cmx150μm, 1.5μm, Bruker), and analyzed on a high-resolution TIMS-TOF Pro2 mass spectrometer (Bruker) using a chromatographic method of 15 “samples per day”. Peptides were ionized by positive-mode electrospray in the captiveSpray source of the TIMS-TOF Pro 2 mass spectrometer, where they were analyzed in DDA (Data-Dependent Acquisition) mode. Thus, each MS scan (between 350 and 1700 Da) was divided into 10 ion mobility ramps (between 0.6 and 1.6 V· s/cm²) and, within each ramp, up to 10 precursors (charge between 2+ and 5+) were selected for fragmentation by High Collision Dissociation (HCD), with the corresponding MS/MS spectra acquired.

Shotgun Protein Identification (FragPipe). The files generated by the DDA analysis were processed offline using the FragPipe pipeline with the human proteome database obtained from the UniProt repository, adding sequences of common contaminants and the reverse database required for validation by false discovery rate (FDR). The default settings of the MSFragger module ^82,83^ were used with some modifications: peptide length from 9 to 30 amino acids, peptide mass range from 300 to 4000, and oxidation of methionine and carbamidomethyl cysteine as modifications (variable and fixed, respectively). In addition, tolerances were set for the precursor (±20 ppm) and fragments (±20 ppm). PSM validation was performed using the Percolator algorithm ^84^, and protein inference was performed using PeptideProphet ^85^, followed by filtering to an FDR of 0.01 ^86^.

Alternatively, in the case of the MAFB immunoprecipitation from resting M-MØ, supernatant from beads treated with Glycine 0.2M pH 2.3 were subjected to SDS-PAGE gel on the Mini-Protean system (BioRad), with a 10% acrylamide acrylamide:bis-acrylamide, 19:1 separating gel approximately 4 cm long in 1.5 M Tris, pH 8.8, and a 4 cm long 4% acrylamide concentrator gel in 0.5 M Tris, pH 6.5. The complete sample, divided into supernatant and magnetic beads, was applied to each well after boiling for 5 minutes in loading buffer (6 mM Tris, 2% SDS, 10% glycerol, 0.5 M β-mercaptoethanol, traces of bromophenol blue) and cooled for 5 minutes at 4°C. Electrophoresis was performed in Laemmlís buffer at 100 V until the electrophoresis front (bromophenol blue) had advanced approximately 2 centimeters in the focusing gel. Protein bands were visualized by staining with colloidal Coomassie (G-250). To do this, the gels were fixed in 50% methanol and 2% phosphoric acid for 30 min. The fixing solution was removed by two washes with milli-Q water, and the gel was equilibrated in 33% methanol, 17% ammonium sulfate, and 3% phosphoric acid for 45 minutes. Afterward, a Coomassie G-250 solution in methanol was added to a concentration of 6.6 mg/ml and left overnight. The next day, the excess Coomassie was removed by washing with milli-Q water for as long as necessary. To carry out the “In-Gel Digestion”, protein bands from the concentrating gel were excised from the gel, and traces of Coomassie were removed by performing two washes with acetonitrile (ACN), alternating with rehydration of the gel with 25 mM ammonium bicarbonate (AMBI). Disulfide bonds were reduced with 10 mM DTT in 25 mM AMBI at 56°C for 30 min, and cysteine thiol groups blocked with 22.5 mM iodoacetamide in 25 mM AMBI for 10 minutes in the dark. After removing residual reagents with two ACN washes, the gel was completely dehydrated in a Speed-Vac (Thermo-Savant) for 30 minutes, after which trypsin (sequencing grade, Roche) was added at a concentration of 12.5 ng/µl in 25 mM AMBI and left to incubate overnight at 37 °C. The peptides were collected from the digestion supernatant, and any remaining peptides in the gel were eluted with ACN. The resulting solution was dried in the Speed-Vac and reconstituted in 20 µL of 2% ACN, 0.1% formic acid.

In this case, DDA analysis proceeded using 20 µL of the digestion product for mass spectrometry analysis. For this purpose, the peptides were injected into the Vanquish Neo nanoHPLC (Thermo) and separated on a 50-cm column (EASY-SPRAY PEPMAP RSLC C18 2UM, 50CMX75UM, Thermo) using an ACN concentration gradient ranging from 2% to 40% over 60 minutes. The chromatographically separated peptides were ionized by positive-mode electrospray and analyzed on a Q Exactive HF mass spectrometer (Thermo) in DDA (data-dependent acquisition) mode. From each MS scan (between 350 and 1700 Da), the 10 most intense precursors (charge between 2+ and 5+) were selected for HCD (high collision energy dissociation) fragmentation, and the corresponding MS/MS spectra were acquired. The data files generated during the shotgun analysis were transferred to the Proteome Discoverer software (Thermo), where the PSMs (peptide-spectrum matches) for each MS/MS spectrum were identified by comparing them with lists of theoretical masses corresponding to the mass of the original precursor, extracted from the human protein database in the UniProt sequence repository using the Sequest search engine. The peptides identified in this manner were assigned to their corresponding proteins. When a peptide could be assigned to multiple proteins, the software uses the principle of parsimony to generate a “master” protein, which is reported in the results. The Percolator algorithm was used to estimate the FDR (false discovery rate), and a q-value <0.01 was applied to filter for proteins identified with high confidence. The mass spectrometry proteomics data have been deposited to the ProteomeXchange Consortium via the PRIDE ^87^ partner repository with the dataset identifier PXD083076.

### Western blot

Cell lysates were subjected to SDS-PAGE (50 μg unless indicated otherwise) and transferred onto an Immobilon-P polyvinylidene difluoride membrane (PVDF; Millipore). After blocking the unoccupied sites with 5% non-fat milk diluted in Tris-Buffered Saline plus Tween 20 (TBS-T), protein detection was carried out with antibodies against MAFB (#HPA005653, Sigma Aldrich), MAF (#sc-518062, Santa Cruz Biotechnology) and Vinculin (#V9131, Sigma-Aldrich) or GAPDH (#sc-32233, Santa Cruz Biotechnology) as protein loading controls. Chemiluminiscence was detected in a Chemidoc Imaging system (BioRad) using SuperSignal™ West Femto (ThermoFisher Scientific).

### RNA-sequencing and data analysis

RNA isolated from M-MØ transfected with siRNA MAF (ΔMAF M-MØ), siRNA MAFB (ΔMAFB M-MØ) or siRNA Control (CNT M-MØ), M-MØ generated in the presence of 1-3 doses of an anti-IL-10 blocking antibody (one dose: M-MØ+anti-IL10*; three doses: M-MØ+anti-IL10) or an isotype-matched control antibody (one dose: M-MØ+IgG*; three doses: M-MØ+IgG), and subjected to sequencing on a BGISEQ-500 platform (https://www.bgitechsolutions.com). Low-quality reads and reads with adaptors or unknown bases were filtered to get the clean reads. Sequences were mapped to GRCh38 genome using HISAT2 ^88^ or Bowtie2 ^89^, and clean reads for each gene were calculated using htseq-count ^90^ and the RSEM software package ^91^. Differential gene expression was assessed by using the R package DESeq2. Differentially expressed genes were analyzed for annotated gene sets enrichment using ENRICHR (http://amp.pharm.mssm.edu/Enrichr/) ^49,92^, and enrichment terms considered significant with a Benjamini-Hochberg adjusted p-value < 0.05. For Gene Set Enrichment Analysis (GSEA) (http://software.broadinstitute.org/gsea/index.jsp) ^51^, gene sets available at the website, as well as gene sets generated from publicly available transcriptional studies (https://www.ncbi.nlm.nih.gov/gds), were used. The datasets used throughout the manuscript (either reported here for the first time or previously published) are listed and described in Supplementary Table S1. RNAseq data were deposited in the Gene Expression Omnibus (http://www.ncbi.nlm.nih.gov/geo/) under accession GSE205719 (M-MØ transfected with siRNA specific for MAF, MAFB or Control, and either untreated or exposed to CL264) and GSE344310 (M-MØ generated in the presence of anti-IL-10 or an isotype-matched antibody).

### Cleavage Under Targets and Tagmentation (CUT&Tag)

CUT&Tag was performed using the Hyperactive Universal CUT&Tag Assay Kit for Illumina (TD904, Vazyme) in combination with the Hyperactive TF Enhancer Module for CUT&Tag (TD904-C2, Vazyme), according to the manufacturer’s instructions. For each condition, 200.000 human M-MØ were used as input, and immunoprecipitation performed with a primary antibody against MAFB (HPA005653, Sigma-Aldrich), MAF (#26745, Cell Signaling Technology), or no-antibody control. Raw paired-end reads were adapter-and quality-trimmed with fastp v1.1.0 ^93^ using automatic paired-end adapter detection. Reads were required to have a Phred quality score ≥ 20, with no more than 30% of bases falling below this threshold and one ambiguous base (N) at most per read. Reads shorter than 25 bp after trimming were discarded. Trimmed reads were aligned to the human reference genome GRCh38.p14, using Bowtie2 v2.5.4 ^89^ with parameters optimized for transcription-factor CUT&Tag (--very-sensitive-local, --no-unal, --maxins 1500). Aligned fragments were filtered to retain confidently mapped reads (MAPQ ≥ 30), coordinate-sorted, and indexed with SAMtools v1.22.1^94^. Reads overlapping ENCODE hg38 blacklist regions ^95^ were removed using BEDTools v2.31.1 ^96^. Genome-wide coverage tracks were generated from the filtered alignments with bamCoverage from the deepTools v3.5.6 suite ^97^ at single-base resolution (--binSize 1), with reads centered and extended to their full fragment length, and normalized to counts per million mapped reads (CPM). Peaks were called with MACS2 v2.2.9.1 ^98^ in paired-end mode (-f BAMPE) against the human effective genome size. As CUT&Tag produces sparse background, model building was disabled and fragments were assigned a fixed extension of 200 bp (--nomodel --shift 0 -- extsize 200), all duplicate reads were retained (--keep-dup all). Enriched regions were called at a q-value threshold of 0.01, and signal-per-million-reads−normalized bedGraph tracks were produced (--SPMR --bdg). Reproducible binding sites were defined per condition using DiffBind v3.20.0 ^99^. For each condition (MAFB, MAF, no-antibody), a consensus peakset was constructed from the MACS2 peak calls and retained only those intervals detected in all three replicates of the condition, yielding a high-confidence set of reproducible peaks that were exported for downstream analysis. Consensus peaks were annotated to genomic features with ChIPseeker v1.46.1 ^100^ and HOMER’s annotatePeaks.pl ^101^ against the UCSC hg38 known-gene models, with the promoter region defined as-1000/+100 bp around the transcription start site (TSS). CUT&Tag data were deposited in the Gene Expression Omnibus (http://www.ncbi.nlm.nih.gov/geo/) under accession GSE345993.

### Phagocytosis assay

Phagocytosis ability was assessed by flow cytometry using pHrodo Red *E. coli* BioParticles Conjugates (Thermo Fisher), following the procedures recommended by the manufacturer. Macrophages were cultured in 24-well plates and exposed to pHrodo bioparticles for 60 min at 37 °C and 5% CO2. Cells were then harvested and assessed by flow cytometry.

### Efferocytosis assay

Jurkat cells were cultured in RPMI 1640 medium without FBS for 16h, and then treated with staurosporine (0.5 μg/ml, Sigma), followed by an incubation at 37 °C for 3h. Staurosporine treatment yielded a population of 73% annexin V + cells. Apoptotic cells (AC) were resuspended at a concentration of 1 × 10^6^ cells/ml and labeled with CellTrace™ Violet reagent (0.5 μM, Invitrogen, Thermo Scientific) for 20 min. For efferocytosis, macrophages were cultured with labeled AC (ratio 1:4) in p12 plates during 1h at 37 °C. Then, macrophages were rinsed with PBS to remove unbound AC and detached with PBS 10 mM EDTA, and pelleted by centrifugation before analyzing by flow cytometry.

### Allogeneic Mixed Lymphocyte Reaction

Naïve CD4+ T cells were isolated from human PBMCs (from donors different from those used for macrophage generation) using the Naïve CD4+ T Cell Isolation Kit (Miltenyi Biotec) in an autoMACS Pro separator (Miltenyi Biotec), according to the manufacturer’s instructions. Purified naïve CD4+ T cells were subsequently labeled with 5 µM CFSE (Invitrogen). Macrophages were detached using PBS-EDTA, collected, counted and resuspended in complete RPMI 1640 medium [RPMI 1640, Lonza, supplemented with 10% heat-inactivated fetal bovine serum (FBS, HyClone), 100 µg/mL Normocin (InvivoGen), 50 µg/mL penicillin-streptomycin, 1% nonessential amino acids, 1% MEM vitamins and 1 mM sodium pyruvate (all from Life Technologies)]. For MLR, 1 × 10^5^ macrophages were cocultured with 5 × 10^5^ naïve CD4+ T cells (1:5 macrophage-to-T-cell ratio) in 48-well plates in a final volume of 500 µL per well. Cocultures were maintained for 5 days at 37°C in a humidified atmosphere containing 5% CO3. Following the 5-day coculture, cells and supernatants were collected for downstream analyses. Cells were stained for viability using Aqua (Invitrogen), followed by surface staining with anti-CD4 (BD Biosciences), anti-CD25 (BioLegend), and anti-CD127 (BioLegend). Cells were subsequently fixed and permeabilized using FOXP3 Fix/Perm and Perm/Wash Buffers (BioLegend) and stained intracellularly with anti-FOXP3-Alexa Fluor 647 (BioLegend). Flow cytometry analysis was performed to assess T-cell activation and proliferation based on CFSE dilution, with particular emphasis on proliferation within the CD4+CD25+CD127−FOXP3+ regulatory T-cell (Treg) population.

### Definition of the Training Signature Score

The “Training Signature Score” was calculated based on a reference set of approximately 500 genes characterized by increased H3K4me3 enrichment at their regulatory regions and transcriptional upregulation in β-glucan-trained human monocytes ^72,74^. To provide a physiological baseline the reference scores for “Naive” and “Trained” states were derived from a consensus β-glucan training dataset ^72,74^. Therefore, this score is a quantitative measure to determine how much a specific condition (e.g., MAF knockdown) “mimics” the epigenetic and transcriptional state of a β-glucan-trained monocyte. Specifically, the “Trained Reference Gene Set” was generated with the genes that are transcriptionally upregulated (Fold Change>2; FDR<0.05) by β-glucan, exhibit LPS-responsiveness, and show a significant H3K4me3 in monocytes 6 days after β-glucan exposure ^72,74^. This “Trained Reference Gene Set” is dominated by genes encoding pro-inflammatory cytokines (*TNF, IL6, IL1B, IL12B*), chemokines (*CCL2, CCL5, CXCL8, CXCL10*), metabolic enzymes (*HK2, PFKP, LDHA, SLC2A1*), surface markers / receptors (*CD80, CD86, TLR4, PLA2G2A*) and other genes involved in transcription / signaling (*MYC, HIF1A, IRF1, NAMPT*). The Training Signature Score was calculated using z-score transformation (normalization of the expression levels and z-score Calculation for every gene in the “Trained Reference Gene Set” in ΔMAF M-MØ and ΔMAFB M-MØ), Thus, a Score of 0 means the knockdown has no effect on trained immunity genes, whereas a score of 2.24 (e.g., in the case of ΔMAF M-MØ) means that, on average, the genes that define trained immunity are shifted more than 2 standard deviations higher.

The Score essentially represents the mean Log2 Fold Change (LFC) of the exclusive regulatory modules for MAF and MAFB that overlap with the core TLR4/7-inducible program. While ΔMAF M-MØ yielded a score of 2.24, ΔMAFB M-MØ only reached 0.67 is due to the coverage of the genes affected. This is so because MAF knockdown derepresses a massive number of inflammatory cytokines and chemokines that are “primed” during β-glucan training, causing the average score to be very high. In fact, the high “Training Signature Score” of ΔMAF M-MØ results from a high connectivity (nearly 80% of these genes are upregulated simultaneously, leading to a very high mean z-score) and the magnitude of the change (with genes like *IL1B* and *CXCL10* showing a >10-fold increase upon MAF silencing), effectively recapitulating the trained state in the absence of exogenous priming. By contrast, MAFB knockdown primarily derepresses the metabolic portion of the trained signature (*HK2*, *PKM*), which represent a smaller percentage of the total “Trained Reference Gene Set”, thus resulting in a lower “Training Signature Score” (0.67).

### Statistical analysis

Statistical analyses were conducted using the GraphPad Prism software. For comparison of means, and unless otherwise indicated, statistical significance of the generated data was evaluated using the paired Student t test or One-Way ANOVA followed by multiple comparisons using the Dunnett’s test or Fisheŕs LSD test. In all cases, p<0.05 was considered as statistically significant.

## DATA AVAILABILITY

The data set supporting the conclusions of this article is available in the Gene Expression Omnibus repository (http://www.ncbi.nlm.nih.gov/geo/) under accessions GSE205719, GSE344310, GSE345993 (CUT&TAG for MAFB or MAF). The mass spectrometry proteomics data have been deposited to the ProteomeXchange Consortium via the PRIDE partner repository with the dataset identifier PXD083076

## ACKNOWLEDGEMENTS

This work was supported by: grants PID2020-114323RB-I00 and PID2024-157082OB-I00 (to ALC), and grant PID2024-156862OB-I00 (to OP) from Ministerio de Ciencia, Innovación y Universidades/ Agencia Estatal de investigación (MICIU/AEI), and co-funded by “FEDER: A way to make Europe”; and grant P2022/BMD-7274 (to ALC) from Dirección General de Innovación e Investigación Tecnológica de la Comunidad de Madrid. ACB, IR, MSM and ACJ were funded by a Formación de Personal Investigador predoctoral fellowships from Ministerio de Ciencia e Innovación (Grants PRE2022-101330, PRE2021-097080, PRE2018-083396 and PREP2024-002306, respectively).

## AUTHOR CONTRIBUTIONS

Conceptualization: A.L.C., M.A.V., O.P.

Methodology: A.C.B., I.R., M.S.F., A.C.J., C.H., A.M., I.Z., F.C., A.A.

Data analysis: A.C.B., I.R., M.S.F., A.C.J., C.H., A.M., I.Z., F.C., A.A., O.P., M.A.V., A.L.C.

Writing, original draft: A.L.C.; Writing, review & editing, A.L.C., M.A.V., O.P., A.C.B., I.R.

Funding acquisition & Resources: A.L.C., M.A.V., O.P.

Supervision: A.L.C.

## CONSENT TO PARTICIPATE DECLARATION

Not Applicable

