## Supplemental Tables for "MAFB and MAF cooperatively orchestrate human anti-inflammatory macrophage identity via autocrine Interleukin-10"

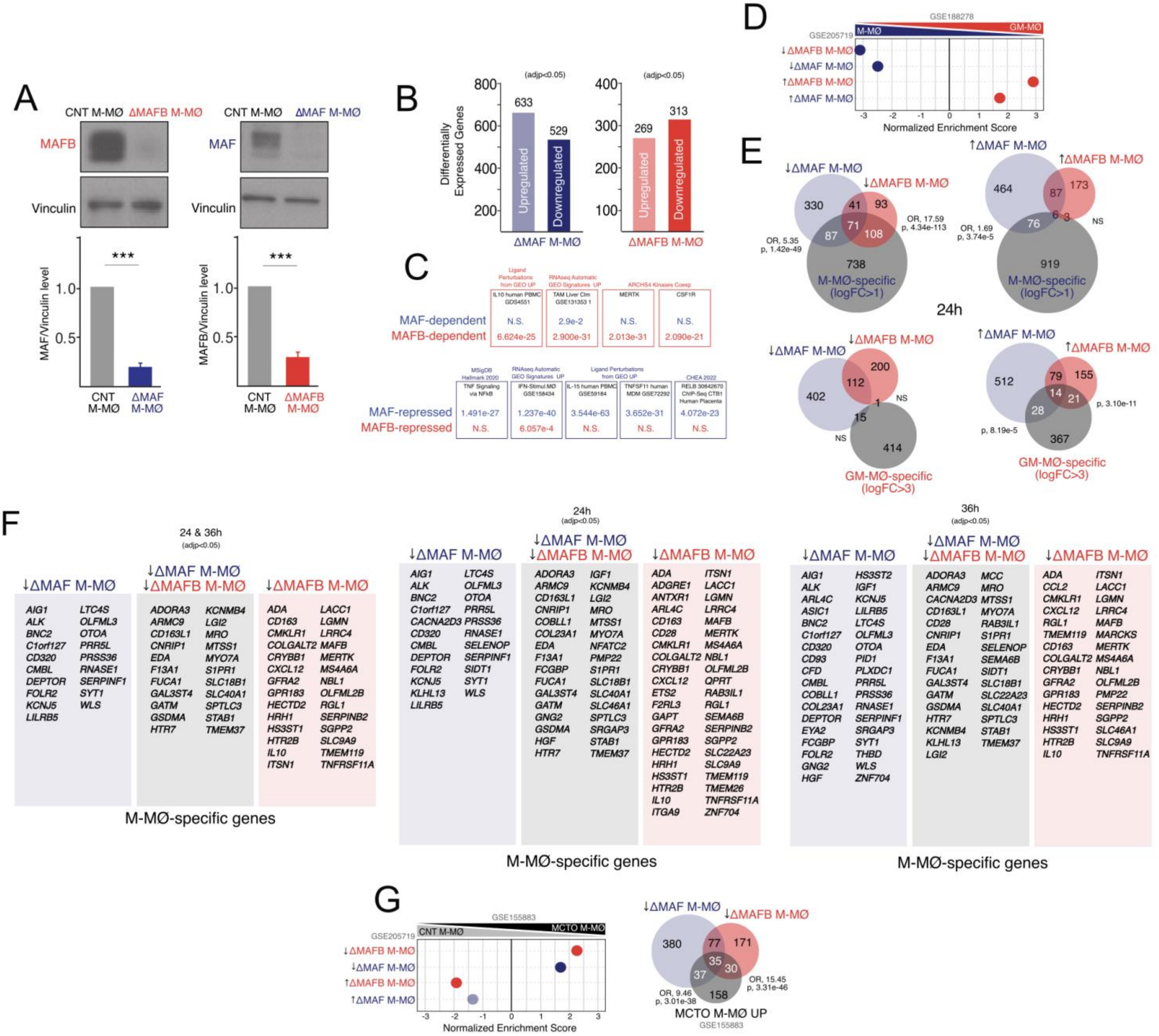

Supplementary Figure 1

**Supplementary Figure 1. A.** Western blot validation showing selective knockdown of MAF and MAFB protein in ΔMAF M-MØ and ΔMAFB M-MØ, respectively. Vinculin protein levels were determined as protein loading control. Mean ± SEM of the MAFB/Vinculin and MAF/Vinculin protein ratios from three independent experiments are shown (*Lower panel*) (paired Student's t-test: ***, p<0.005). Representative western blot experiments are shown (*Upper panel*). **B**. Number of differentially expressed genes (adjp<0.05) between ΔMAF M-MØ (or ΔMAFB M-MØ) and CNT M-MØ. Differential gene expression was assessed using DESeq2. **C**. Gene ontology analysis of MAFB- and MAF-regulated gene sets using Enrichr databases. **D**. Summary of GSEA of MAF- and MAFB-regulated gene sets (GSE205719) on the ranked comparison of the transcriptomes of M-MØ vs. GM-MØ (GSE188278). NES and FDRq values are indicated in each case (Solid-colored dots, FDRq=0). **E**. Venn diagram overlaps (Hypergeometric test) of the gene sets defining the core M-MØ-specific (logFC>1) or GM-MØ-specific (logFC>3) identity and the genes downregulated or upregulated in ΔMAF M-MØ or ΔMAFB M-MØ at 24h after silencing. **F**. Identification of M-MØ-specific genes (logFC>3) significantly (adjp<0.05) downregulated in ΔMAF M-MØ, ΔMAFB M-MØ, or ΔMAF M-MØ and ΔMAFB M-MØ, at the indicated times after silencing. **G**. Summary of GSEA of MAF- and MAFB-regulated gene sets (GSE205719) on the ranked comparison of the transcriptomes of MCTO M-MØ vs. CNT M-MØ (GSE155883). NES and FDRq values are indicated in each case (Solid-colored dots, FDRq=0; Light-colored dots, FDRq<0.250) (*Upper panel*). Venn diagram overlaps (Hypergeometric test) of the genes upregulated in MCTO M-MØ (MCTO M-MØ UP) and the gene sets downregulated in ΔMAF M-MØ or ΔMAFB M-MØ at 24h after silencing.

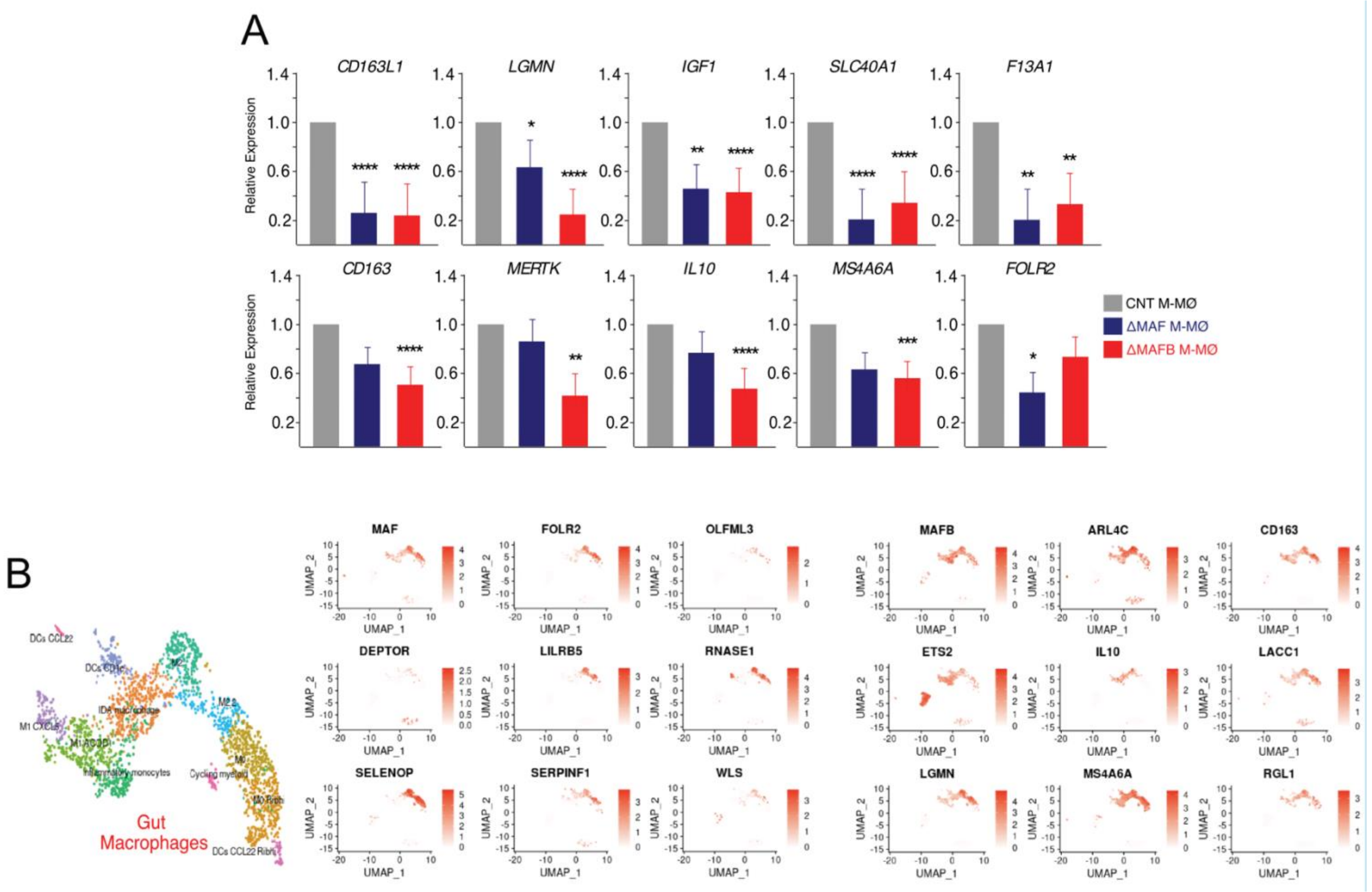


Supplementary Figure 2

**Supplementary Figure 2. A.** Relative mRNA levels of the indicated genes in ΔMAF M-MØ, ΔMAFB M-MØ and CNT M-MØ, as determined by qRT-PCR. Mean ± SEM of three independent samples is shown (paired Student´s t test: *, p <0.05; **, p<0.005; ***, p <0.001). **B.** Color-coded expression of the indicated genes after projection onto the UMAP (Uniform Manifold Approximation and Projection) embedding shown the gut myeloid cell clusters identified in GSE214695 [1].

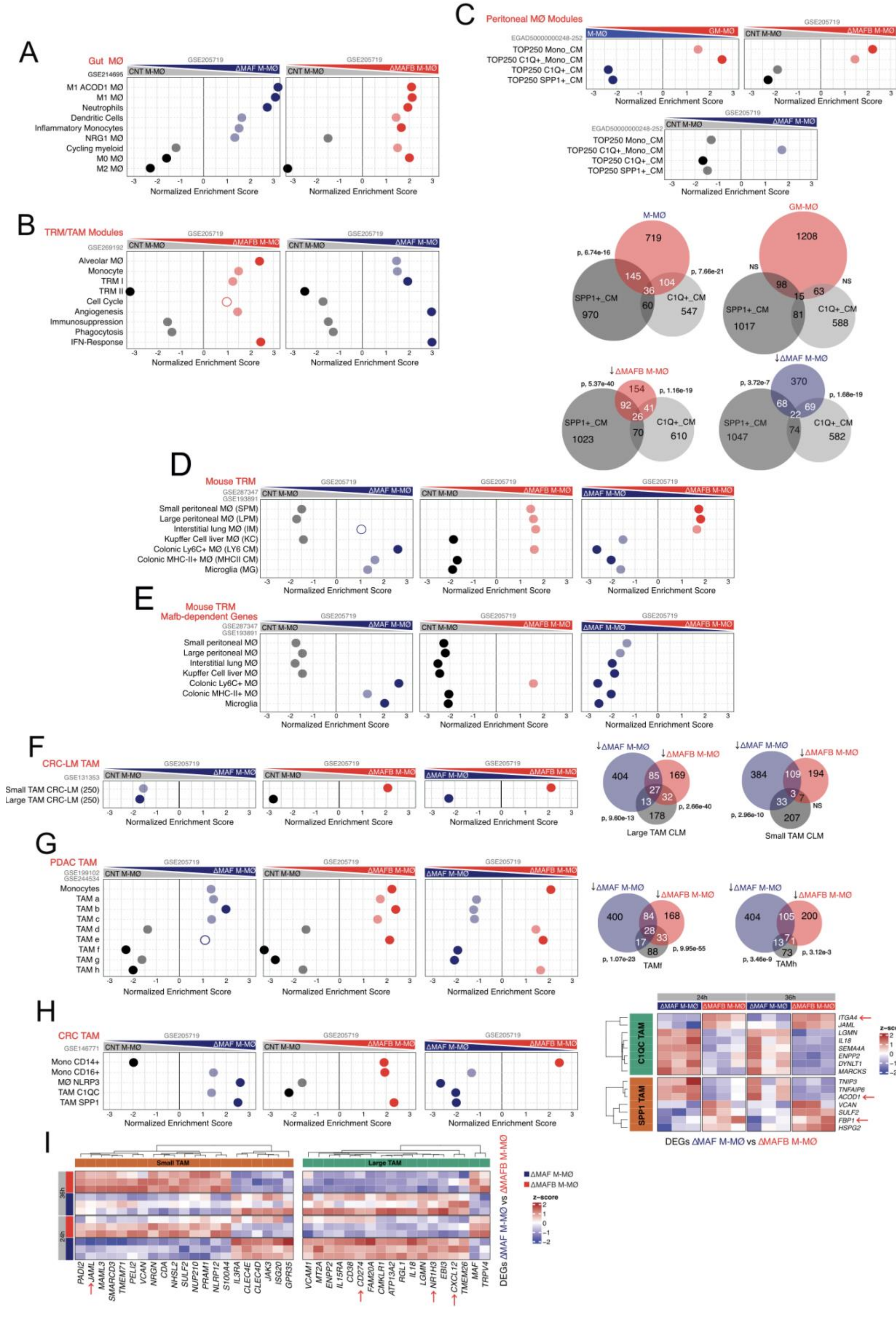


Supplementary Figure 3

**Supplementary Figure 3. A-H.** Summary of GSEA of the gene sets defining myeloid/monocyte/macrophages subsets in various homeostatic and pathological tissues (**A**, Gut macrophages and myeloid cells; **B**, TRM/TAM modules from various tissues; **C**, Peritoneal MØ modules; **D**, Mouse TRM from various tissues; **E**, Mafb-dependent genes within mouse TRM from various tissues; **F**, Colorectal cancer liver metastasis TAM (CRC-LM TAM); **G**, Pancreatic ductal adenocarcinoma TAM (PDAC TAM); **H**, Colorectal cancer TAM (CRC TAM)) on the ranked comparison of ΔMAF M-MØ vs CNT M-MØ, ΔMAFB M-MØ vs CNT M-MØ and ΔMAF M-MØ vs ΔMAFB M-MØ. NES and FDRq values are indicated in each case (Solid-colored dots, FDRq=0; Light-colored dots, FDRq<0.250; Empty dots, Non significant FDRq). In **C**, **F** and **G**, Venn diagram overlaps (Hypergeometric test) of the indicated gene sets are shown. In **H**, heatmap illustrates the preferential expression of the genes differentially expressed between ΔMAF M-MØ and ΔMAFB M-MØ (at 24 h and 36 h post-silencing) in SPP1 TAM or C1QC TAM [2]. **I**. Heatmap highlighting the preferential expression of the genes differentially expressed between ΔMAF M-MØ and ΔMAFB M-MØ (at 24 h and 36 h post-silencing) in Small TAM CRC-LM or Large TAM CRC-LM [3].

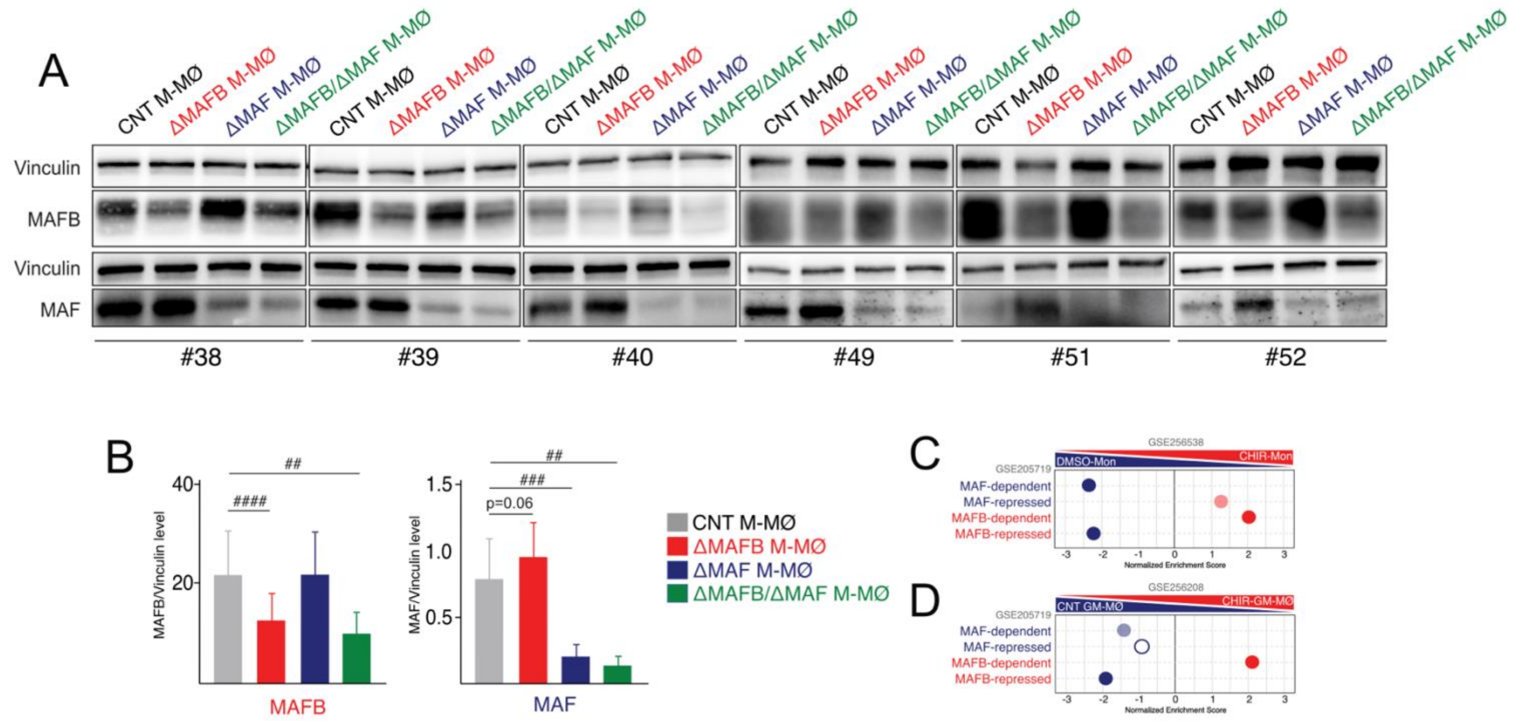


Supplementary Figure 4

**Supplementary Figure 4. A.** Expression of MAF and MAFB protein in six independent samples of ΔMAF M-MØ, ΔMAFB M-MØ, ΔMAF/ΔMAFB M-MØ and CNT M-MØ, as determined by Western blot. Vinculin protein levels were determined as protein loading control. **B**. Relative expression of MAF and MAFB protein in ΔMAF M-MØ, ΔMAFB M-MØ, ΔMAF/ΔMAFB M-MØ and CNT M-MØ. Mean ± SEM of the MAFB/Vinculin and MAF/Vinculin protein ratios from six independent experiments are shown (Ratio-Paired Student's t-test: #, p <0.05; ##, p<0.005; ###, p <0.001). **C-D.** Summary of GSEA of the indicated MAF- or MAFB-regulated gene sets on the ranked comparison of the transcriptomes of (**C**) CHIR-treated monocytes (CHIR-Mon) vs DMSO-Mon or (**D**) CHIR-treated GM-MØ (CHIR-GM-MØ) vs. CNT GM-MØ. NES and FDRq values are indicated in each case (Solid-colored dots, FDRq=0; Light-colored dots, FDRq<0.250; Empty dots, Non significant FDRq).

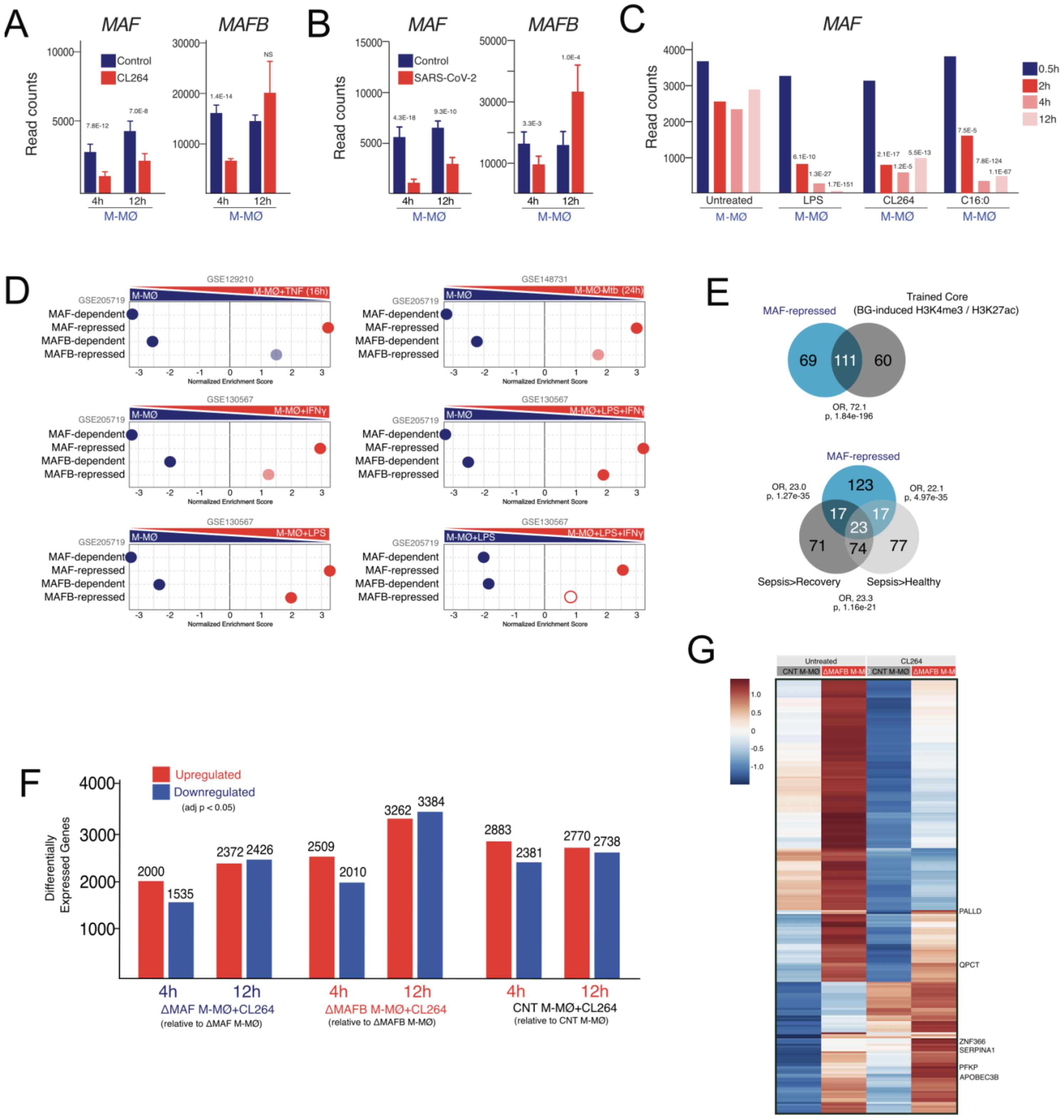


Supplementary Figure 5

**Supplementary Figure 5. A.** *MAF* and *MAFB* expression in M-MØ and M-MØ treated with CL264 for 4h-12h, as determined by RNA-Seq (GSE205719) [45]. Adjp values relative to untreated M-MØ are indicated. **B**. *MAF* and *MAFB* expression in control and SARS-CoV-2-treated M-MØ (4h, 12h), as determined by RNA-Seq (GSE207840) [4]. Adjp values relative to untreated M-MØ are indicated. **C**. *MAF* expression in control and LPS-, CL264- or Palmitate (C16:0)-treated M-MØ at the indicated times after stimulation, as determined by RNA-Seq (GSE156921)[5]. Adjp values relative to untreated M-MØ are indicated. **D**. Summary of GSEA of the indicated gene sets on the ranked comparison of the transcriptomes of M-MØ+TNF(16h) vs. M-MØ (GSE129210), M-MØ+IFNγ vs. M-MØ (GSE130567), M-MØ+LPS vs. M-MØ (GSE130567), M-MØ+LPS+IFNγ vs. M-MØ (GSE130567), M-MØ+*Mtb*(24h) vs. M-MØ (GSE148731), and M-MØ+LPS+IFNγ vs. M-MØ+LPS (GSE130567). NES and FDRq values are indicated in each case (Solid-colored dots, FDRq=0; Light-colored dots, FDRq<0.250; Empty dots, Non significant FDRq). **E**. Venn diagram overlaps (Hypergeometric test) of the MAF-repressed gene set and (*Upper panel*) the "Trained core" gene set derived from datasets in [6,7] or gene sets specifically upregulated in hyper-inflammatory sepsis monocytes [8]. **F**. Number of Differentially Expressed Genes (DEG, adjp<0.05) between the transcriptomes of ΔMAF M-MØ+CL264, ΔMAFB M-MØ+CL264, and CNT M-MØ+CL264, and their corresponding untreated counterparts. **G**. Heatmap of the expression of genes exhibiting an additive or synergistic response to MAFB knockdown and exposure to CL264. Selected genes are indicated.

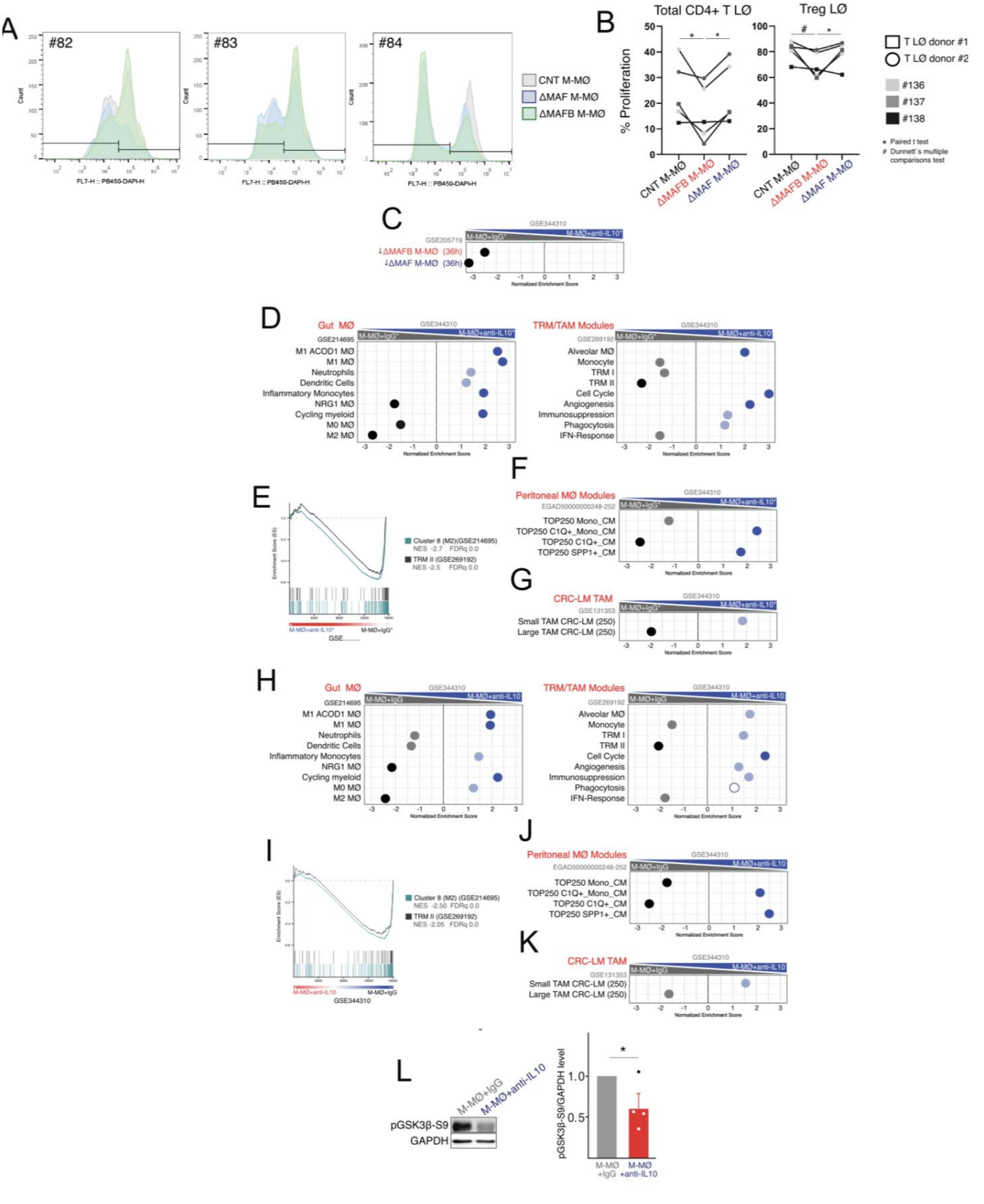


Supplementary Figure 6

**Supplementary Figure 6. A.** Efferocytosis capacity of three independent samples of ΔMAF M-MØ, ΔMAFB M-MØ and CNT M-MØ, as determined by flow cytometry using staurosporine-induced CellTrace Violet-labeled apoptotic Jurkat cells. **B.** Individual results of the five allogeneic MLR promoted by CNT M-MØ, ΔMAF M-MØ or ΔMAFB M-MØ, as determined by flow cytometry on CFSE (carboxyfluorescein succinimidyl ester)-labeled cells to track T cell division. Statistical analysis was performed using a paired Student's t-test (*p < 0.05) for the indicated comparisons and repeated-measures one-way ANOVA with Dunnett's multiple-comparison test (*, p < 0.05). **C.** GSEA of the indicated gene sets on the ranked comparison of the M-MØ+IgG* vs. M-MØ+anti-IL10* transcriptomes. **D-G.** Summary of GSEA of the indicated gene sets on the ranked comparison of the transcriptomes of M-MØ+IgG* vs. M-MØ+anti-IL10* (GSE344310). The specific GSEA profiles for Gut M2 MØ and TRMII are shown in (**E**). **H-K**. Summary of GSEA of the indicated gene sets on the ranked comparison of the transcriptomes of M-MØ+IgG vs. M-MØ+anti-IL10 (GSE344310). The specific GSEA profiles for Gut M2 MØ and TRMII are shown in (**I**). NES and FDRq values are indicated in all GSEA results (Solid-colored dots, FDRq=0; Light-colored dots, FDRq<0.250; Empty dots, Non significant FDRq). **L.** (*Left panel*) Phosphorylated Ser9-GSK3β levels in M-MØ+IgG and M-MØ+anti-IL10, as determined by Western blot. GAPDH protein levels were determined as protein loading control. (*Right panel*) Mean ± SEM of the Ser9-GSK3β/GAPDH protein ratios from four independent experiments are shown (paired Student's t-test: *, p<0.05).
